# The SPPL3-defined glycosphingolipid repertoire regulates immune responses by improving HLA class I access

**DOI:** 10.1101/2020.09.26.313585

**Authors:** Marlieke L.M. Jongsma, Matthijs Raaben, Antonius A. de Waard, Tao Zhang, Birol Cabukusta, René Platzer, Vincent A. Blomen, Anastasia Xagara, Tamara Verkerk, Sophie Bliss, Lennert Janssen, Elmer Stickel, Stephanie Holst, Rosina Plomp, Arend Mulder, Soldano Ferrone, Frans H.J. Claas, Mirjam H.M. Heemskerk, Marieke Griffioen, Hermen Overkleeft, Johannes B. Huppa, Manfred Wuhrer, Thijn R. Brummelkamp, Jacques Neefjes, Robbert M. Spaapen

**Affiliations:** Department of Immunopathology, Sanquin Research, Amsterdam, The Netherlands; Landsteiner Laboratory, Amsterdam UMC, University of Amsterdam, Amsterdam, The Netherlands; Oncode Institute and Department of Cell and Chemical Biology, LUMC, Leiden, The Netherlands; Department of Biochemistry, The Netherlands Cancer Institute, Amsterdam, The Netherlands; Center for Proteomics and Metabolics, LUMC, Leiden, The Netherlands; Institut für Hygiene und Angewandte Immunologie, Vienna, Austria; Department of Cell Biology II, The Netherlands Cancer Institute, Amsterdam, The Netherlands; Department of Immunohaematology and Blood Transfusion, LUMC, Leiden, The Netherlands; Department of Surgery, Massachusetts General Hospital, Harvard Medical School, Boston, MA, United States; Department of Hematology, Leiden University Medical Center, Leiden, The Netherlands; Leiden Institute of Chemistry, Leiden University, Leiden, The Netherlands; CeMM Research Center for Molecular Medicine of the Austrian Academy of Sciences, Vienna, Austria; CGC.nl, Amsterdam, The Netherlands.

**Keywords:** HLA class I, glycosphingolipids, B3GNT5, SPPL3, T cell activation, antigen presentation

## Abstract

HLA class I (HLA-I) drives immune responses by presenting antigen-derived peptides to cognate CD8^+^ T cells. This process is often hijacked by tumors and pathogens for immune evasion. Since therapeutic options for restoring HLA-I antigen presentation are limited, we aimed to identify new HLA-I pathway targets. By iterative genome-wide screens we uncovered that the cell surface glycosphingolipid (GSL) repertoire determines effective HLA-I antigen presentation. We show that absence of the protease SPPL3 augments B3GNT5 enzyme activity, resulting in upregulated levels of surface (neo)lacto-series GSLs. These GSLs sterically impede molecular interactions with HLA-I and diminish CD8^+^ T cell activation. In accordance, a disturbed SPPL3-B3GNT5 pathway in glioma associates with decreased patient survival. Importantly, we show that this immunomodulatory effect can be reversed through GSL synthesis inhibition using clinically approved drugs. Overall, our study identifies a GSL signature that functionally inhibits antigen presentation and represents a potential therapeutic target in cancer, infection and autoimmunity.

## Introduction

Human Leukocyte Antigen class I (HLA-I) proteins are the primary modules recognized by CD8^+^ T cells determining both the induction and effector phase of immune responses. Their primary function is to present peptide fragments from degraded proteins to the T cell receptor (TCR) of cytotoxic CD8^+^ T cells, leading to T cell activation (Neefjes et al., 2011; Unanue and Cerottini, 1989). Because HLA-I molecules on tumor cells also present tumor antigen-derived peptides to cognate T cells, they play a major role in the anti-tumor activity of T cells unleashed by current immunotherapeutic strategies (Schumacher and Schreiber, 2015). As a consequence, tumors often escape from these therapies by downregulating one or multiple molecules critical in HLA-I antigen presentation (Chowell et al., 2018; Gettinger et al., 2017; Restifo et al., 1996; Sade-Feldman et al., 2017; Zaretsky et al., 2016). This reduction is often reversible, for example by interferon stimulation, ionizing radiation or inhibition of histone deacetylases, which has led to various experimental therapies aimed at increasing tumor HLA-I surface expression (Haworth et al., 2015; Reits et al., 2006; Thor Straten and Garrido, 2016). Moreover, sensitizing tumor cells as immune targets can act synergistically with T cell (re)activating strategies, thereby increasing the therapeutic potential of enhancing HLA-I availability (Hahnel et al., 2008). Active suppression of HLA-I surface expression to escape T cell surveillance is also employed by pathogens, such as Epstein-Barr virus and cytomegalovirus (Hansen and Bouvier, 2009; Yewdell and Hill, 2002). These and other examples underscore the broad relevance of HLA-I based interventions, necessitating a thorough understanding of the molecular mechanisms underpinning the HLA-I pathway.

Over the last 35 years, several key elements in HLA-I expression and antigen presentation have been identified and extensively studied. A protein complex directed by the transcriptional regulator NLRC5 drives HLA-I expression in selected tissues while HLA-I translation and glycosylation are currently thought to be executed by general enzymes and mechanisms (Jongsma et al., 2017; Ryan and Cobb, 2012). Once synthesized, the HLA-I heavy chain and its light chain beta-2 microglobulin (B2M) assemble and are stabilized by a unique combination of the ER chaperone proteins tapasin, ERp57 and calreticulin (CALR) (Rock et al., 2016). Two of these HLA-I chaperone complexes bind the peptide transporter TAP to form the so-called peptide loading complex (PLC), which drives efficient ER import and loading of peptides into the HLA-I peptide-binding groove (Blees et al., 2017). Subsequently, mature trimeric complexes of HLA-I heavy chain, B2M and peptide are released from the PLC for transport to the cell surface for peptide presentation to T cells (Garstka et al., 2015; Wearsch and Cresswell, 2008). Given the multifactorial complexity of the HLA-I antigen presentation pathway, we hypothesized that additional regulatory mechanisms of this central process in adaptive immunity must exist.

To uncover new components of the HLA-I antigen presentation pathway we carried out a series of genome-wide haploid genetic screens. In addition to the known factors described above, we identified the enzyme signal peptide peptidase like 3 (SPPL3) as a major player in HLA-I antigen presentation. We found that SPPL3 controls the composition of the cell surface glycosphingolipid (GSL) repertoire by inhibiting the glycosyltransferase B3GNT5. In the absence of SPPL3, an increase in B3GNT5 activity leads to high levels of complex negatively charged (neo)lacto-series GSLs (nsGSLs), interfering with activation of CD8^+^ T cells and preventing HLA-I from being accessed by its natural ligand LIR-1. Furthermore, we show that the nsGSL synthesis pathway negatively affects survival prognosis in glioma and that FDA-approved drug intervention with the synthesis of GSLs in glioma model cells results in improved anti-tumor immunity. Hence, the cellular GSL composition represents a novel and targetable layer of immune regulation by determining the quality of antigen presentation.

## Results

### A haploid genetic screen provides a high-resolution map of the HLA-I antigen presentation pathway

To identify unknown factors regulating HLA-I antigen presentation, we performed a genome-wide insertional mutagenesis screen in haploid human HAP1 cells endogenously expressing HLA-I (Brockmann et al., 2017; Carette et al., 2009). A heterogeneous pool of millions of cells, each knockout (KO) for a random gene or set of genes, was generated by retroviral gene trap insertion and expanded. Mutagenized cells that were poorest or best recognized by the HLA-I-specific W6/32 antibody were sorted by flow cytometry (Figure 1A). Subsequently, we determined the relative enrichment of unique disruptive integrations per gene between both sorted populations based on deep sequencing. This provided an unbiased overview of genes involved in HLA-I antigen presentation at an unprecedented resolution (Figure 1B). Among the highly significant positive modifiers were the known key genes directing HLA-I expression, such as the transcriptional (co-)activators NLRC5, RFXAP and RFX5 (Jongsma et al., 2017) and essential components for post-transcriptional assembly, glycosylation and peptide loading (B2M, MOGS (α-glucosidase I), GANAB (α-glucosidase II), TAP1, TAP2, tapasin, ERp57 and CALR) (Figure 1B) (Wearsch and Cresswell, 2008). Thus, this single genetic map identified the components in the HLA-I pathway previously discovered through decades of research. In addition, HLA-I regulators identified in human B cell lymphoma CRISPR/Cas9 screens by Dersh et al. were also highly significant in our haploid genetic screen, such as SUSD6, SND1, ANKRD33B, EZH2 (Dersh et al., submitted). The most prominent hit from our screen in the W6/32^low^ sorted population was the gene encoding SPPL3, a protease that had never been described before in the context of antigen presentation (Figure 1B). SPPL3 is an ER- and Golgi-localized transmembrane protein of the family of intramembrane-cleaving aspartyl proteases (Fluhrer and Haass, 2007).

**Figure 1.**
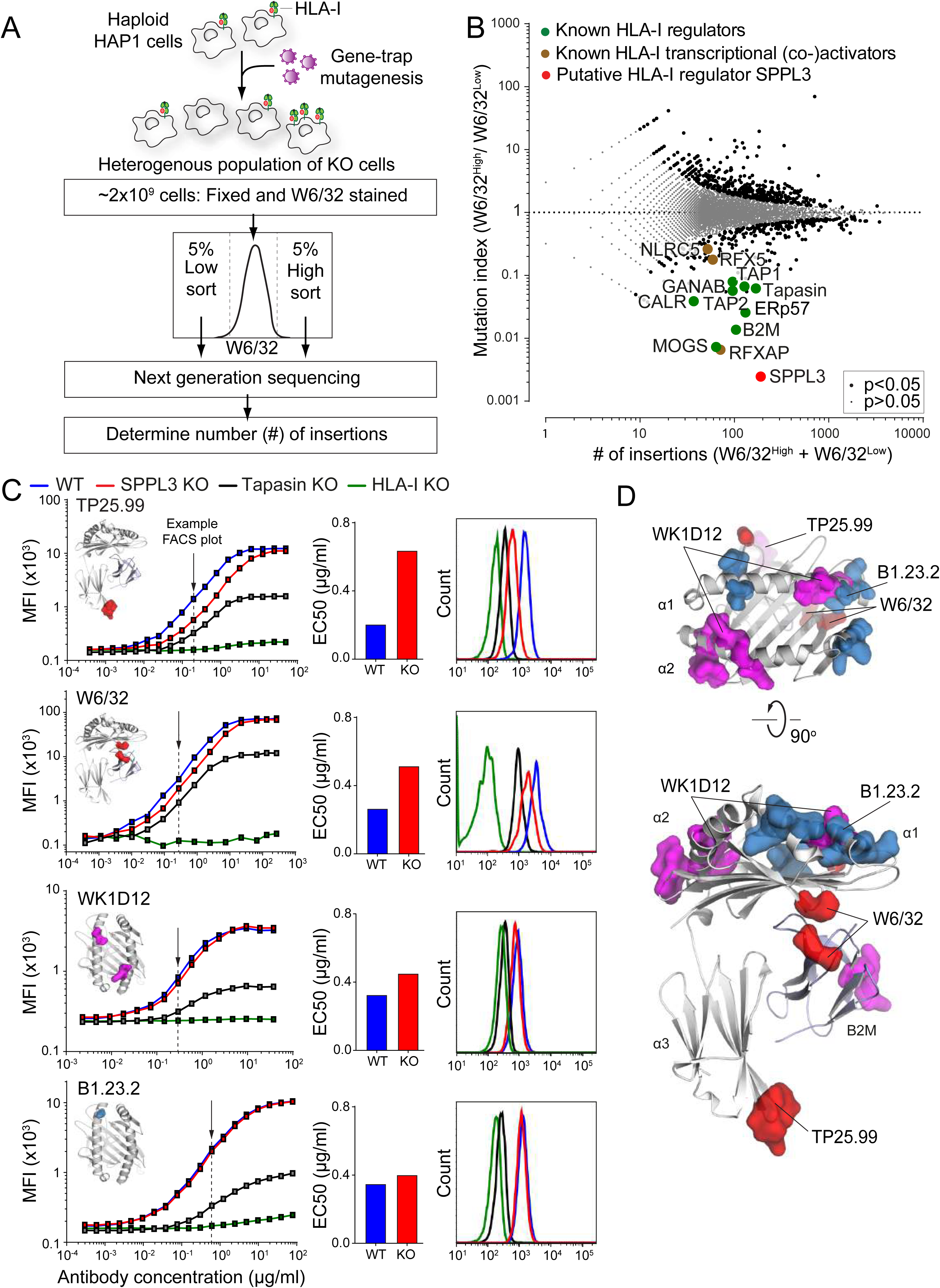
A haploid genetic screen reveals SPPL3 as a novel regulator of accessibility to membrane proximal HLA-I regions. (A) Schematic overview of the genome-wide haploid genetic screen. Human haploid HAP1 cells were mutagenized by retroviral gene trap insertion to create random genetic knockouts. The cells were stained using the W6/32 antibody that recognizes the HLA- A, -B and -C alleles on HAP1 cells and the 5% lowest and 5% highest fluorescent populations were FACS sorted. Sorted populations were sequenced to reveal unique and disruptive gene trap integration sites. (B) A fish-tail plot showing the mutation index (the ratio of integrations mapped per gene in the W6/32^High^ / W6/32^Low^ sorted populations) against the total amount of mapped integrations per gene (W6/32^High^ + W6/32^Low^). Positive and negative regulators of HLA-I (*black/color*) were identified by two-sided Fisher’s exact test, FDR (Benjamini-Hochberg) corrected p<0.05. Known HLA-I regulators are depicted in green, known HLA-I transcriptional (co-)activators in brown and the novel regulator SPPL3 in red. (C) (*left*) Representative titration curves of W6/32, TP25.99, B1.23.2 and WK1D12 antibodies on mixed barcoded (see Figure S1C) HAP1 WT (*blue*), SPPL3 KO (*red*), tapasin KO (*black*) and HLA-I KO cells (*gray*) (see Table S1). The individual antibody binding epitopes are depicted on the HLA-I structure. (*middle*) Titration based EC50 values for WT and SPPL3 KO titrations. (*right*) Histograms of non-saturating antibody stain (concentration chosen around EC50 value as indicated by the arrow) on HAP1 WT (*blue*), SPPL3 KO (*red*), tapasin KO (*black*) and HLA-I KO cells (*gray filled*). n=3 (D) Crystal structure of HLA-I / B2M. SPPL3-susceptible epitopes (*red*), mildly affected epitopes (*purple*) and SPPL3-independent epitopes (*blue*) are highlighted on the structure (see Figure S1D for individual epitopes).

### SPPL3 determines the accessibility of membrane proximal regions of HLA-I

To validate that HLA-I cell surface expression was altered by SPPL3, we created SPPL3 KO cells using CRISPR/Cas9 (Table S1) and performed flow cytometry using the W6/32 antibody. HLA-I surface levels in the absence of SPPL3 turned out identical to those of wild type (WT) cells (Figure S1A), an unexpected result in view of the screening data. Likewise, the total HLA-I content of SPPL3 KO and WT cells was similar as determined by Western blot analysis (Figure S1B). As the anti-HLA antibody W6/32 was used under non-saturating staining conditions in the genome-wide screen, these seemingly contradictory outcomes may have resulted from a reduced accessibility of the W6/32 epitope in the absence of SPPL3.

To test this hypothesis, we titrated the W6/32 antibody for binding WT, SPPL3 KO and control HLA-I KO and tapasin KO cells (Table S1), which were individually color-barcoded and mixed in a single well for optimal comparison of staining intensity (Figure S1C). In contrast to saturating W6/32 concentrations, lower W6/32 concentrations resulted in decreased binding to SPPL3 KO cells compared to WT cells, indicating that access of the HLA-I epitope recognized by W6/32 was indeed hindered (Figure 1C). A similar result was apparent using monovalent W6/32 Fab fragments, validating this conclusion (Figure S1D). To further define SPPL3-dependent HLA-I accessibility, we performed additional titrations using 14 HLA-I-specific antibodies recognizing distinct HLA-I epitopes and one targeting a B2M epitope. The binding of three antibodies (clones W6/32, TP25.99, ROU9A6) was markedly affected by the absence of SPPL3 (Figures 1C and S1D). By superimposing critical amino acid positions for binding of the individual antibodies onto an HLA-I structure, we defined that the SPPL3-susceptible region is relatively proximal to the cellular membrane, an area which is largely conserved among HLA-I alleles (Figure 1D). Binding of several antibodies (e.g. B1.23.2) was not affected by SPPL3, further supporting that HLA-I surface levels are not targeted by SPPL3 and providing unique intra-molecular controls for further experiments. To determine whether SPPL3 differentially affected HLA-A, -B or -C alleles, we reconstituted HLA-A, -B and -C KO cells on a WT or SPPL3 KO background (Table S1) with the single original HLA-I alleles and analyzed their accessibility. Each allele showed a comparable difference in HLA-I accessibility between WT and SPPL3-deficient cells as detected by W6/32 (Figure S2A). In addition, other cell lines exhibited a similar decrease in HLA-I accessibility after siRNA knockdown or CRISPR/Cas9 KO of SPPL3, indicating that regulation of HLA-I accessibility is not solely restricted to HAP1 cells or their HLA-I haplotype (Figures S2B, S2C and S2D).

### SPPL3 expression promotes HLA-I ligand binding and CD8^+^ T cell activation

The fact that SPPL3 modulates antibody reactivity towards specific regions of HLA-I molecules raises the question whether the HLA-I function is affected. The defined SPPL3-susceptible region is the binding site of the inhibitory receptor LIR-1, which is expressed by monocytes, B cells and some subsets of other immune cells (Borges et al., 1997; Colonna et al., 1997) (Figure 2A). Even more pronounced than for SPPL3-affected antibodies, binding of a recombinant LIR-1 Fc fusion protein (Gonen-Gross et al., 2010) to HLA-I on SPPL3 KO cells was strongly decreased compared to WT cells, suggesting that immune cell function can be largely affected by SPPL3 (Figure 2B). A second immune cell receptor ligating closely to the SPPL3-affected HLA-I region is the CD8 costimulatory molecule on T cells (Figure 2A), which in most cases is essential for effective docking of a TCR to its cognate peptide HLA-I complex and sensitizing T cell recognition (Gao et al., 1997; Purbhoo et al., 2004; Roszkowski et al., 2003). We evaluated whether SPPL3 enhances HLA-I antigen presentation to CD8^+^ T cells by stimulating multiple HLA-A*02:01-restricted T cell clones specific for different tumor-expressed antigens with WT and SPPL3 KO cells (Amir et al., 2011; van Bergen et al., 2007; Van Bergen et al., 2010). All clones were more reactive to SPPL3 expressing WT cells, as determined by their IFN-γ or GM-CSF production (Figure 2C), indicating that SPPL3 increases functional access to HLA-I.

**Figure 2.**
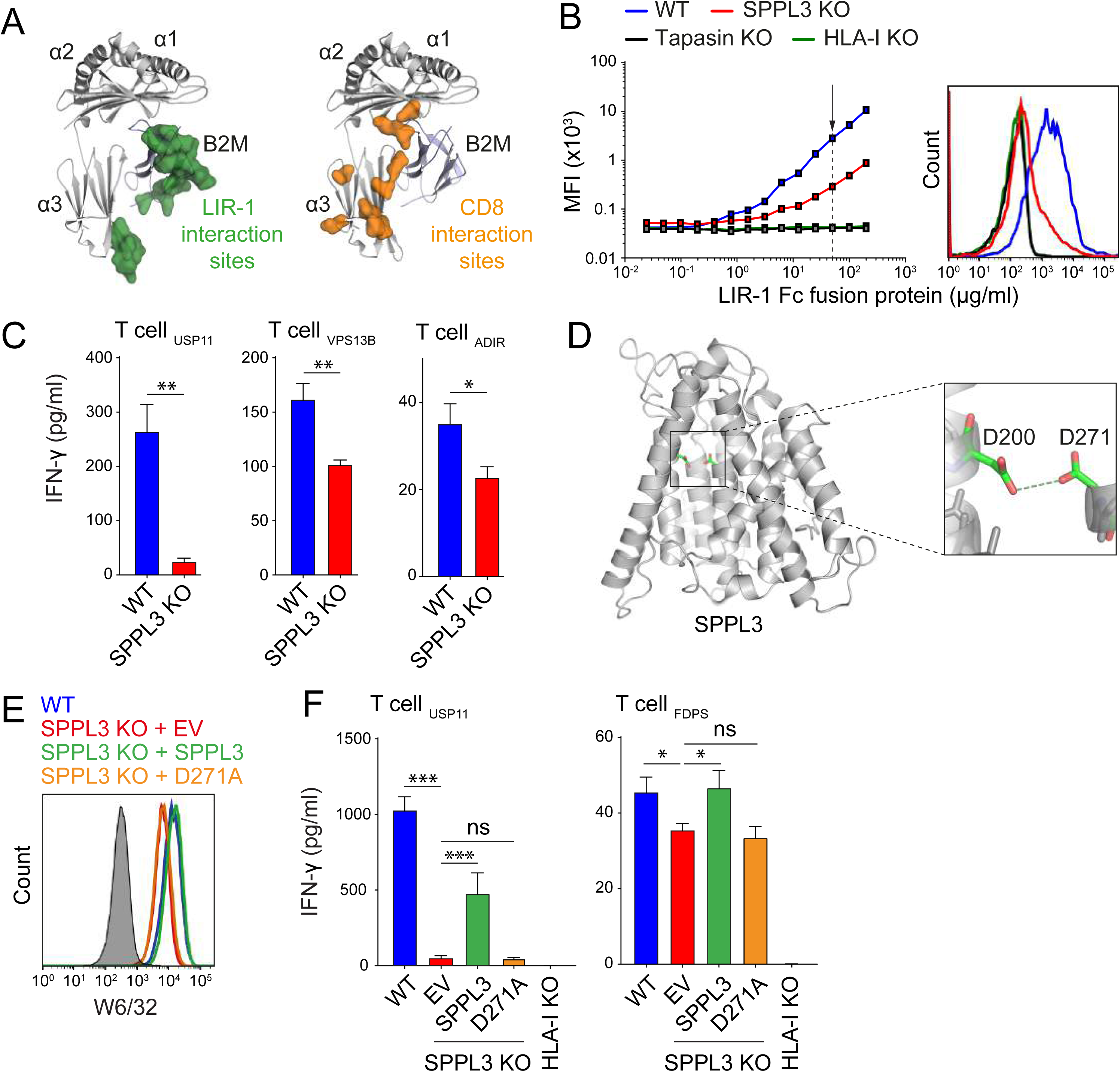
SPPL3 expression promotes LIR-1 binding to HLA-I and enhances CD8^+^ T cell activation. (A) LIR-1 interaction sites (*left, green*) and CD8 interactions sites (*right, orange*) mapped on the crystal structure of HLA-I / B2M. (B) (*left*) Representative titration curves of LIR-1 Fc fusion protein on HAP1 WT (*blue*), SPPL3 KO (*red*), tapasin KO (*black*) and HLA-I KO cells (*green*). (*right*) Representative histogram of LIR-1 Fc binding from the indicated concentration (arrow) on HAP1 WT (*blue*), SPPL3 KO (*red*), tapasin KO (*black*) and HLA-I KO cells (*green*). n=2 (C) IFN-γ production by HLA-A*0201-restricted T cells recognizing endogenously derived USP11, VPS13B and ADIR peptides after overnight co-culture with HAP1 WT (*blue*) or SPPL3 KO (*red*) cells determined by ELISA. Representative of n=3 (D) Predicted protein structure of SPPL3 with its catalytic residues magnified in the detailed view. (E) Representative histogram of non-saturating W6/32 stain on HAP1 WT (*blue*) or SPPL3 KO cells transduced with either RFP-empty vector (*EV, red*), RFP-SPPL3 (*green*) or catalytically inactive RFP-SPPL3 D271A (*orange*). Unstained control is in gray. Transduced cells shown are within RFP^+^ gate. See Figure S2E for SPPL3-independent B1.23.2 antibody stain. n=2 (F) IFN-γ secretion by HLA-A*0201-restricted T cells recognizing endogenously presented USP11 or FDPS peptides after overnight co-culture with HAP1 SPPL3 KO cells transduced with either RFP- empty vector (*EV, red*), RFP-SPPL3 (*green*) or catalytically inactive RFP-SPPL3 D271A (*orange*) followed by FACS sort enrichment for RFP positive cells or WT (*blue*) or HLA-I KO (*gray*) cells, as determined by ELISA. n=1

### HLA-I accessibility depends on SPPL3-mediated proteolytic cleavage of a novel target

SPPL3 contains two aspartate residues embedded in conserved YD and GxGD motifs located in transmembrane helices 6 and 7 respectively forming the catalytic unit for proteolysis (Voss et al., 2013) (Figure 2D). To investigate whether SPPL3 catalytic activity was critical in controlling HLA-I accessibility and function, we expressed wild type SPPL3 or a catalytically inactive SPPL3 mutant (D271A) in SPPL3 KO cells (Voss et al., 2012). Flow cytometry analysis showed that only SPPL3 D271A failed to restore the accessibility of HLA-I for W6/32, indicating that SPPL3 proteolytic activity is required for antibody access to HLA-I (Figure 2E and S2E). This lack of rescue was further confirmed on a functional level since expression of active but not inactive SPPL3 D271A in SPPL3 KO cells partially restored their capacity to activate T cells (Figure 2F).

SPPL3 has previously been reported to affect protein *N*-glycosylation by proteolytic inactivation of glycosyltransferases in the ER and Golgi (Kuhn et al., 2015; Voss et al., 2014). Variations in the *N*- linked glycan of HLA-I, located at position N86 in close proximity to the SPPL3-susceptible region (Figure S2F), can affect its accessibility (Barbosa et al., 1987; Neefjes et al., 1990). However, liquid chromatography-mass spectrometry (LC-MS) of HLA-I *N*-linked glycans revealed no differences between the profiles of WT and SPPL3 KO cells (Figure S2G), indicating that the HLA-I *N*-glycan structure is not regulated by SPPL3 activity. To further exclude a contribution of the HLA-I *N*-glycan to SPPL3-modulated HLA-I accessibility, we inhibited complex *N*-glycan formation on SPPL3 KO cells using the α-mannosidase I and II inhibitors kifunensine and swainsonine. The decrease of complex *N*- glycans failed to alter the accessibility of the W6/32 epitope on SPPL3 KO cells (Figures S2H and S2I). This result was confirmed in cells genetically engineered to lack complex (HLA-I) *N*-glycosylation through ablation of the gene encoding GANAB (Table S1), which resulted in lower overall HLA-I surface levels as visualized by decreased W6/32 and B1.23.2 signals (Figure S2J). Comparison of these antibody stainings between WT and SPPL3 KO cells showed that the HLA-I accessibility was still impaired (Figure S2J). As we ruled out a role for protein glycosylation, our finding that SPPL3 activity affects HLA-I at the cell surface suggests the involvement of at least one currently unknown SPPL3 target.

### SPPL3-controlled glycosphingolipids modulate HLA-I accessibility

To elucidate how SPPL3 controls HLA-I accessibility, we followed two genome-wide screening strategies to specifically identify targets that are either positively or negatively regulated by SPPL3. An SPPL3-activated target affecting HLA-I accessibility would likely be a hit in the original W6/32 screen, just like SPPL3 (Figure 1B). However, the identification of such a target was complicated by the long list of significant hits. To distinguish SPPL3-activated targets from other candidates, we complemented the original screen with a new genome-wide haploid screen using a different HLA-I-specific antibody that was significantly less affected by the absence of SPPL3 (antibody BB7.2) (Figures 3A and S1D). This additional screen yielded another high-resolution snapshot of HLA-I antigen presentation (Figure S3A). A comparison of the outcome of the two screens showed that SPPL3 was the only factor selectively affecting W6/32 binding, implying that no other gene was as strongly required for the accessibility of HLA-I (Figure 3B).

**Figure 3.**
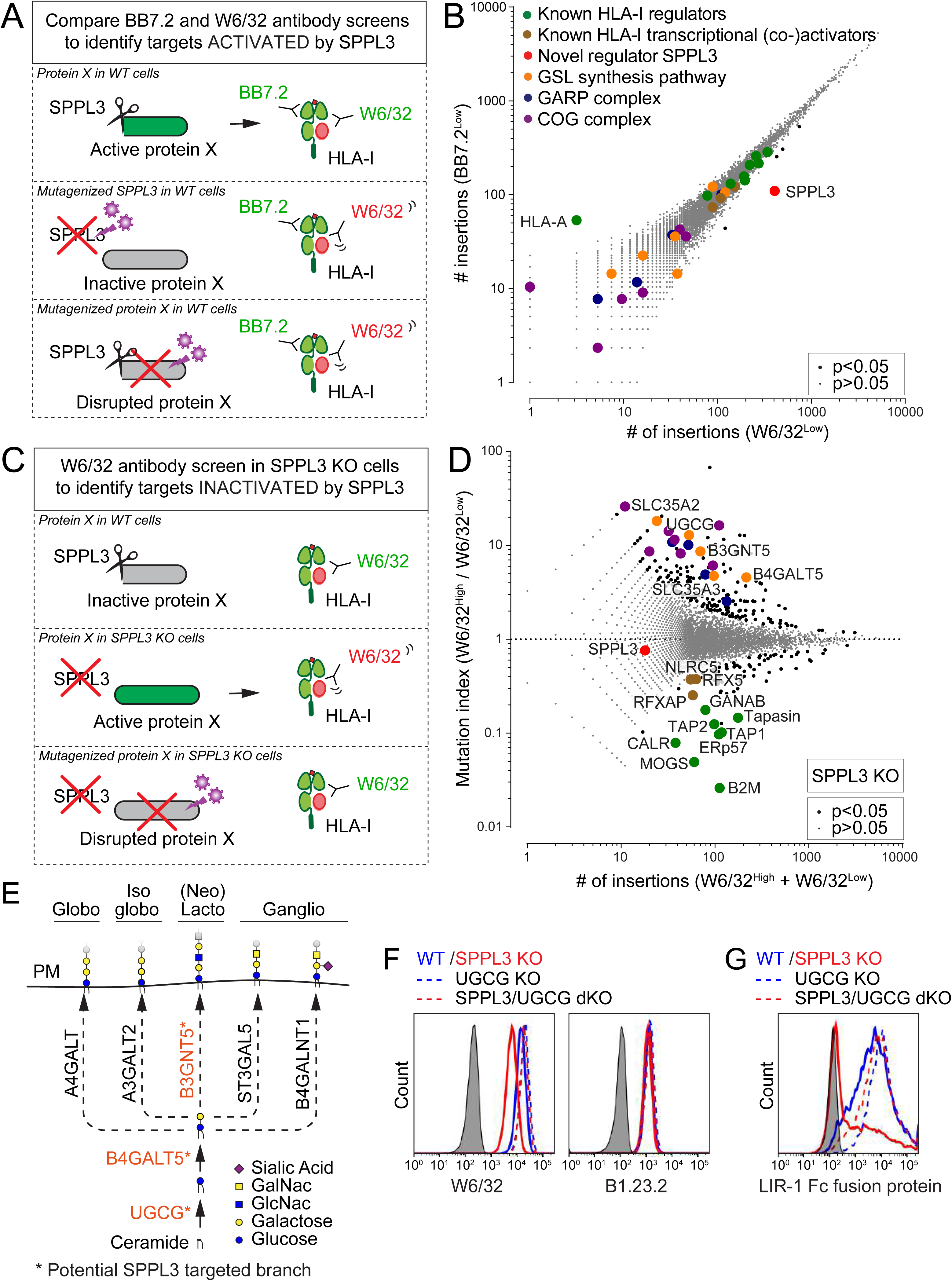
SPPL3-controlled glycosphingolipids modulate accessibility of HLA-I. (A) Schematic outline showing that a combination of BB7.2 and W6/32 antibody screens will lead to the specific identification of potential HLA-I regulators activated by SPPL3. (B) Rocket plot depicts the number of unique disruptive integrations per gene in the BB7.2Low sorted population plotted against the number of unique disruptive integrations per gene in the W6/32^Low^ sorted population. Positive and negative regulators of HLA-I (*black/color*) were determined by Fisher’s exact test, FDR (Benjamini-Hochberg) corrected p<0.05. Highlighted are known HLA-I regulators (*green*), HLA-I transcriptional (co-)activators (*brown*), the regulator SPPL3 (*red*), proteins involved in the glycosphingolipid (GSL) synthesis pathway (*orange*) and members of the GARP and COG complexes (*blue and purple respectively*). (See also Figure S3A). (C) Schematic outline showing that a W6/32 antibody screen in SPPL3 KO cells will lead to the identification of potential HLA-I regulators that are normally inactivated by SPPL3 activity in WT cells. (D) Fish-tail plot of the mutation index (ratio of integrations per gene of W6/32^High^ / W6/32^Low^ populations) against the total amount of integrations per gene (W6/32^High^ + W6/32^Low)^. Positive and negative regulators of HLA-I (*black/color*) were determined by Fisher’s exact test, FDR (Benjamini-Hochberg) corrected p<0.05. Legend as in (B). (See also Figure S3B) (E) Schematic overview of the GSL synthesis pathway. The core enzymes UGCG and B4GALT5 catalyze the first two steps of GSL synthesis. Five branching enzymes initiate the synthesis of four different groups of GSLs: globo-, isoglobo-, (neo)lacto- and ganglio-series. The putative SPPL3-targeted branch is shown in orange. PM, plasmamembrane. (F) Representative histograms of non-saturating W6/32 and B1.23.2 cell surface staining of HAP1 WT (*blue*), SPPL3 KO (*red*), UGCG KO (*blue dashe*d) and SPPL3/UGCG double KO (dKO) cells (*red dashed*). Unstained control is in gray. n=3 (G) Representative histogram of LIR-1 Fc fusion protein binding to HAP1 WT (*blue*), SPPL3 KO (*red*), UGCG KO (*blue dashe*d) or SPPL3/UGCG double KO (dKO) cells (*red dashed*). Unstained control is in gray. n=3

We then searched for potential genes negatively regulated by SPPL3 to affect HLA-I. To this end, we performed a genome-wide haploid screen in SPPL3 KO HAP1 cells. In these cells, which potentially lacked SPPL3-mediated suppression of the sought target, gene trap mutagenesis of such a target or its associated pathway should improve W6/32 access to HLA-I (Figure 3C). The hits from this screen converged to the glycosphingolipid (GSL) synthesis pathway (Figure 3D). The enzymes UGCG, B4GALT5 and B3GNT5 catalyze the synthesis of GSLs in the Golgi membrane by consecutive linkage of sugar residues derived from activated UDP-glucose, UDP-galactose and UDP-*N*- acetylglucosamine donors on ceramide molecules (Figure 3E) (Allende and Proia, 2014). The latter two carbohydrate donors are transported from the cytoplasm into the Golgi by SLC35A2 and SLC35A3 respectively, which were also identified in the screen (Figure 3D) (Caffaro and Hirschberg, 2006). Other hits from the screen included proteins and complexes associated with Golgi homeostasis such as the members of the component of oligomeric Golgi (COG) and Golgi associated retrograde protein (GARP) complexes that further facilitate GSL synthesis and trafficking (Frohlich et al., 2015; Kingsley et al., 1986) (Figure 3D). Of note, none of these hits related to GSL metabolism emerged in the original screen with W6/32 on SPPL3-containing WT cells, strongly suggesting that in WT cells the GSL synthesis or transport pathway is suppressed by SPPL3 (Figure S3B). Collectively, these observations revealed the existence of a pathway comprising GSL-mediated regulation of HLA-I access and function controlled by SPPL3.

To validate that SPPL3 reduces HLA-I accessibility through manipulation of GSL synthesis, we generated GSL-deficient SPPL3 KO cells by additionally knocking out the first enzyme of the GSL synthesis pathway, UGCG (Table S1). Strikingly, in these SPPL3/UGCG double KO cells we observed full rescue of the W6/32 HLA-I epitope accessibility without affecting the SPPL3- independent B1.23.2 staining (Figure 3F) pointing towards an essential role for GSLs in HLA-I accessibility. Importantly, GSL-deficient SPPL3 KO cells regained the capacity to engage the HLA-I ligand LIR-1 (Figure 3G), underscoring the physiological relevance of the effect of GSLs on HLA-I.

### B3GNT5 tunes the capacity of HLA-I to interact with its natural receptors

The synthesis of GSLs is probably best illustrated as a chain of sugar moiety transfers catalyzed by different Golgi enzymes. UGCG initiates the GSL synthesis pathway by transferring a glucose to a ceramide on the cytosolic leaflet of the Golgi membrane (Allende and Proia, 2014). After this glucosylceramide is flipped into the Golgi lumen, a galactose moiety is added by B4GALT5 or B4GALT6 to generate lactosylceramide. This neutral GSL then serves as a substrate for various glycosyltransferases responsible for the generation of different GSL-series: A4GALT (globo-series), A3GALT2 (isoglobo-series), B3GNT5 ((neo)lacto-series; nsGSLs), B4GALNT1 (gangliosides, o-series) and ST3GAL5 (gangliosides, a-,b-,c-series) (Figure 3E) (Allende and Proia, 2014; Zhang et al., 2019). Our screening data suggest that specifically nsGSL production by means of B3GNT5 activity can diminish HLA-I accessibility (Figure 3D). To confirm this specificity, we generated polyclonal cell lines in the SPPL3 KO background, each CRISPR/Cas9-targeting one of the five branching enzymes, and analyzed W6/32 epitope accessibility by flow cytometry. As shown in Figures 4A and S4A, HLA-I accessibility in SPPL3 KO cells was fully restored only by ablation of B3GNT5 or control UGCG, confirming B3GNT5 as the sole branching enzyme involved in HLA-I epitope shielding. This effect was selective for SPPL3-deficient cells as HLA-I accessibility was unaffected on WT cells with corresponding gene KOs (Figure S4B). Single cell derived B3GNT5 KO and SPPL3/B3GNT5 double KO cell lines were generated to further corroborate a pivotal role for B3GNT5 in HLA-I accessibility (Table S1). As expected, additional B3GNT5 KO in SPPL3 KO cells did not only restore W6/32 binding to its epitope, but also the accessibility of other SPPL3-susceptible epitopes recognized by TP25.99 and ROU9A6 (Figure 4B). Accessibility to SPPL3-independent epitopes and total HLA-I surface expression were not affected by additional KO of B3GNT5 (Figure 4B). Most importantly, the lack of B3GNT5 expression in SPPL3 KO cells restored HLA-I functionality by strongly increasing both binding of the ligand LIR-1 as well as the cellular potential to activate T cells (Figures 4C and 4D). Taken together our results suggest that SPPL3 inactivates B3GNT5 through proteolysis to tune the capacity of HLA-I to interact with its receptors.

**Figure 4.**
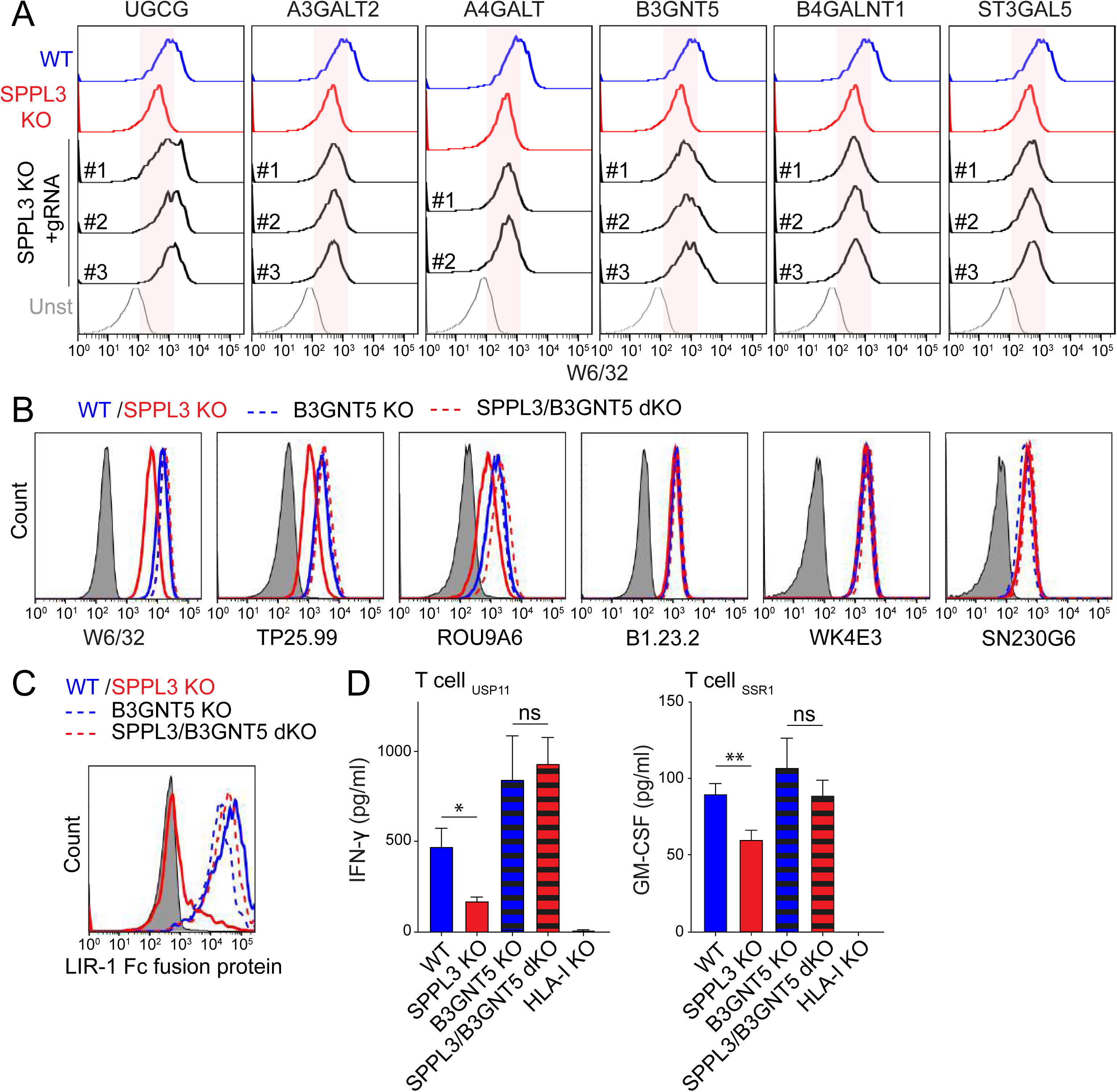
B3GNT5 function determines the HLA-I visibility for its natural receptors. (A) Representative histograms of non-saturating W6/32 cell surface staining of HAP1 WT (*blue*), SPPL3 KO (*red*) or polyclonal populations of SPPL3 KO cells additionally knocked out for the core enzyme UGCG or the branching enzymes A3GALT2, A4GALT, B3GNT5, B4GALNT1 or ST3GAL5 (*black*) by CRISPR/Cas9 using two or three individual guide RNAs (gRNAs, see Table S4) per gene as indicated. Shown are only GFP^+^ transduced cells. The red area indicates fluorescence of SPPL3 KO cells. n=2 (See controls in Figure S5A and S5B) (B) Representative histograms of non-saturating cell surface stainings using three HLA-I specific antibodies recognizing SPPL3-susceptible epitopes (W6/32, TP25.99 and ROU9A6) and three recognizing SPPL3-independent epitopes (SN230G6, WK4E3 and B1.23.2) on HAP1 WT (*blue*), SPPL3 KO (*red*), B3GNT5 KO (*blue dashed*) and SPPL3/B3GNT5 double KO (dKO) cells (*red dashed*). n=3 (C) Representative histograms of LIR-1 Fc fusion protein binding to HAP1 WT (*blue*), SPPL3 KO (*red*), B3GNT5 KO (*blue dashe*d) and SPPL3/B3GNT5 double KO (dKO) cells (*red dashed*). n=2 (D) IFN-γ or GM-CSF secretion by HLA- A*0201-restricted T cells recognizing USP11 or SSR1 peptides after overnight co-culture with HAP1 WT (*blue*), SPPL3 KO (*red*), B3GNT5 KO (*blue dashe*d), SPPL3/B3GNT5 double KO (dKO) (*red dashed*) or HLA-I KO (*gray*) cells was determined by ELISA. n=2 For flow cytometry data, the gray histogram represents an unstained control cell line.

### SPPL3 controls the generation of (neo)lacto-series GSLs by targeting B3GNT5

To visualize a direct interaction between SPPL3 and its putative target B3GNT5, we performed co-immunoprecipitation of overexpressed epitope-tagged proteins. We only co-isolated B3GNT5 with the catalytically inactive SPPL3 D271A mutant suggesting a transient interaction between SPPL3 and its substrate (Figure 5A). Two other branching enzymes of the GSL synthesis pathway, B4GALNT1 and ST3GAL5, were not co-isolated with SPPL3 (Figure 5A). To investigate whether SPPL3 affects B3GNT5 activity, we performed a B3GNT5 enzymatic assay. Lysates of indicated WT and KO cells were incubated with a BODIPY-conjugated analog of the B3GNT5-substrate lactosylceramide (LacCer) and the donor sugar UDP-*N*-acetylglucosamine, followed by thin layer chromatography (TLC) of extracted GSLs. The B3GNT5 product lactotriaosylceramide (BODIPY-Lc3Cer), as confirmed by LC-MS, was generated in increased amounts in SPPL3 KO compared to WT cell lysates (Figures 5B, 5C and S5A). In addition, no Lc3Cer was synthesized in lysates of B3GNT5 KO cells, demonstrating that B3GNT5 is the sole producer of Lc3Cer in HAP1 cells. Since SPPL3 inhibits B3GNT5 activity we next addressed the extent to which SPPL3 defines the cellular GSL profile. Glycan portions of the GSL repertoire of WT, SPPL3 KO and SPPL3/B3GNT5 double KO cells were isolated and analyzed by LC-MS. We found an extensive shift in the relative GSL abundance towards B3GNT5-produced nsGSLs, from 44% in WT cells to 82% in SPPL3 KO cells (Figures 5D, 5E and Table S2). The increase was most evident for complex nsGSLs containing six or more sugar residues as determined by relative quantification of individual GSLs, suggesting that epitope shielding of HLA-I is mediated by complex nsGSLs (Figure S5B and Table S2). To validate the shift in GSL repertoire in living cells, we conducted flow cytometry-based experiments using cholera toxin subunit B, which binds the ganglioside GM1, and an antibody against the nsGSL SSEA-1 epitope. GSL-deficient UGCG KO cells were negative for all probes, demonstrating probe specificity towards GSLs on our cells (Figure S5C). Compared to WT cells, SPPL3 KO cells expressed increased levels of SSEA-1 nsGSLs, which were generated by B3GNT5, and decreased levels of GM1 gangliosides (Figure 5F). Consistent with our relative quantification of individual glycans detected by LC-MS, these data demonstrate that SPPL3 dictates the composition of the GSL repertoire by inhibiting the nsGSL biosynthesis activity of B3GNT5.

**Figure 5.**
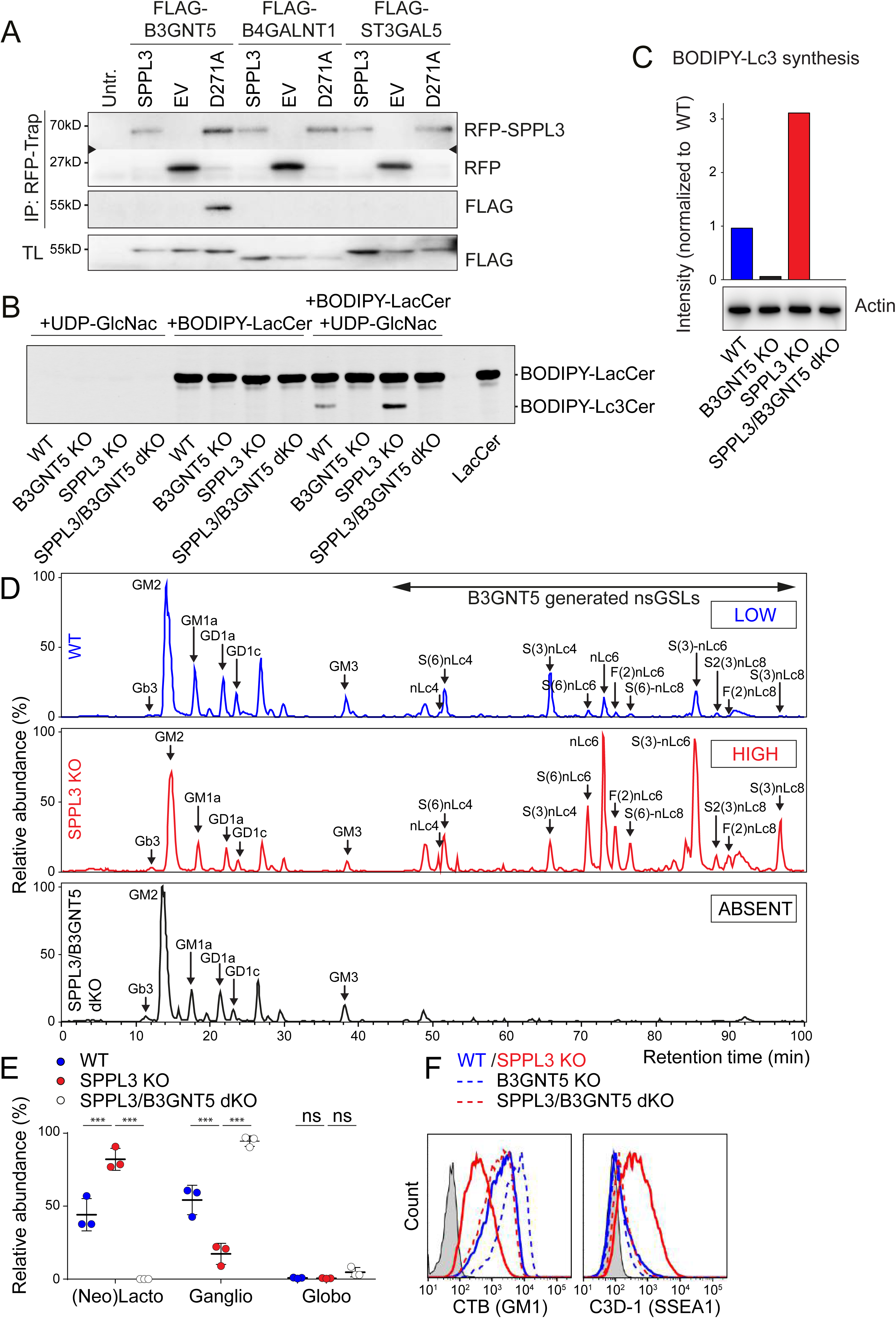
SPPL3 controls the generation of nsGSLs by targeting B3GNT5. (A) Co-immunoprecipitation of RFP-SPPL3, RFP-empty vector (EV) or RFP-SPPL3 D271A with either FLAG- B3GNT5, FLAG-B4GALNT1 or FLAG-ST3GAL5 using RFP-trap beads (intermediate area in the top blot was excised). TL= Total Lysate. n=2 (B) B3GNT5 activity in cell lysates from HAP1 WT, SPPL3 KO, B3GNT5 KO or SPPL3/B3GNT5 double KO (dKO) cells using UDP-GlcNac donor sugar, BODIPY-Lactosylceramide substrate or both, analyzed by thin layer chromatography. n=2 (C) Quantification of B3GNT5 produced BODIPY-Lc3 (see Figure S5A for LC-MS validation). (D) Base peak chromatograms of PGC LC-MS on total GSL glycans isolated from WT (*blue*), SPPL3 KO (*red*) or SPPL3/B3GNT5 double KO (dKO) cells (*black*). The proposed glycan structures, their relative abundance and standard deviation are listed in Table S2 and Figure S6B. Representative of n=3 is shown. (E) Quantified relative abundance of the three subtypes of GSL glycans derived from WT (*blue*), SPPL3 KO (*red*) and SPPL3/B3GNT5 dKO cells (*white*). n=3 (F) Representative histograms of FITC-labeled Cholera Toxin B (anti-GM1) and C3D-1 (anti-SSEA-1) cell surface staining of HAP1 WT (*blue*), SPPL3 KO (*red*), B3GNT5 KO (*blue dashed*) and SPPL3/B3GNT5 double KO (dKO) cells (*red dashed*). Unstained control is in gray. n=3 (See Figure S5C)

### Sialic acid residues on nsGSLs are required for HLA-I shielding

GSLs are major constituents of membrane microdomains (Sezgin et al., 2017). A change in GSL composition may therefore disturb membrane protein localization, mobility and function. We therefore investigated the mobility of HLA-I in SPPL3 KO cells by single particle tracking. The mobile fraction and diffusion constant of BB7.2 Fab labeled HLA-I molecules were equal between SPPL3 KO and WT cells, indicating that HLA-I membrane dynamics were not majorly affected by any potential alterations in membrane microdomain organization nor by SPPL3 itself (Figures S6A, S6B, S6C and S6D). This renders a scenario in which HLA-I associates with another protein in the absence of SPPL3 unlikely, as this would reduce the HLA-I diffusion rate. Instead, our data highly suggest that decreased HLA-I accessibility is a direct consequence of interactions with nsGSLs. Such GSL-protein interactions can occur between gangliosides and hormone receptors through a charge-based linkage of GSL-derived sialic acid with positively charged amino acids (D’Angelo et al., 2013). Further analyses of the GSL signature of SPPL3 KO compared to WT cells revealed that the nsGSL glycan chains more frequently contain α-2,3- and α-2,6-linked sialic acid residues, but also non-charged fucoses (Figures 5D, 6A and Table S2). To test the requirement of these nsGSL-localized sugar residues, we inhibited all sialyl- and fucosyl-transferase activity and found that dose-dependent inhibition of sialylation but not fucosylation restores HLA-I accessibility in SPPL3 KO cells (Figures 6B, 6C, 6D, 6E and 6F). The requirement for sialic acids was further substantiated by the genetic KO of CMP-sialic acid synthetase (CMAS), which recapitulated the recovery of HLA-I accessibility in SPPL3 KO cells (Figure 6G). Finally, the enzymatic removal of sialic acid residues at the cell surface by neuraminidase treatment also diminished HLA-I shielding (Figure 6H). Thus the B3GNT5-generated GSLs shield HLA-I through its sialic acids, probably via a direct charge-based interaction.

**Figure 6.**
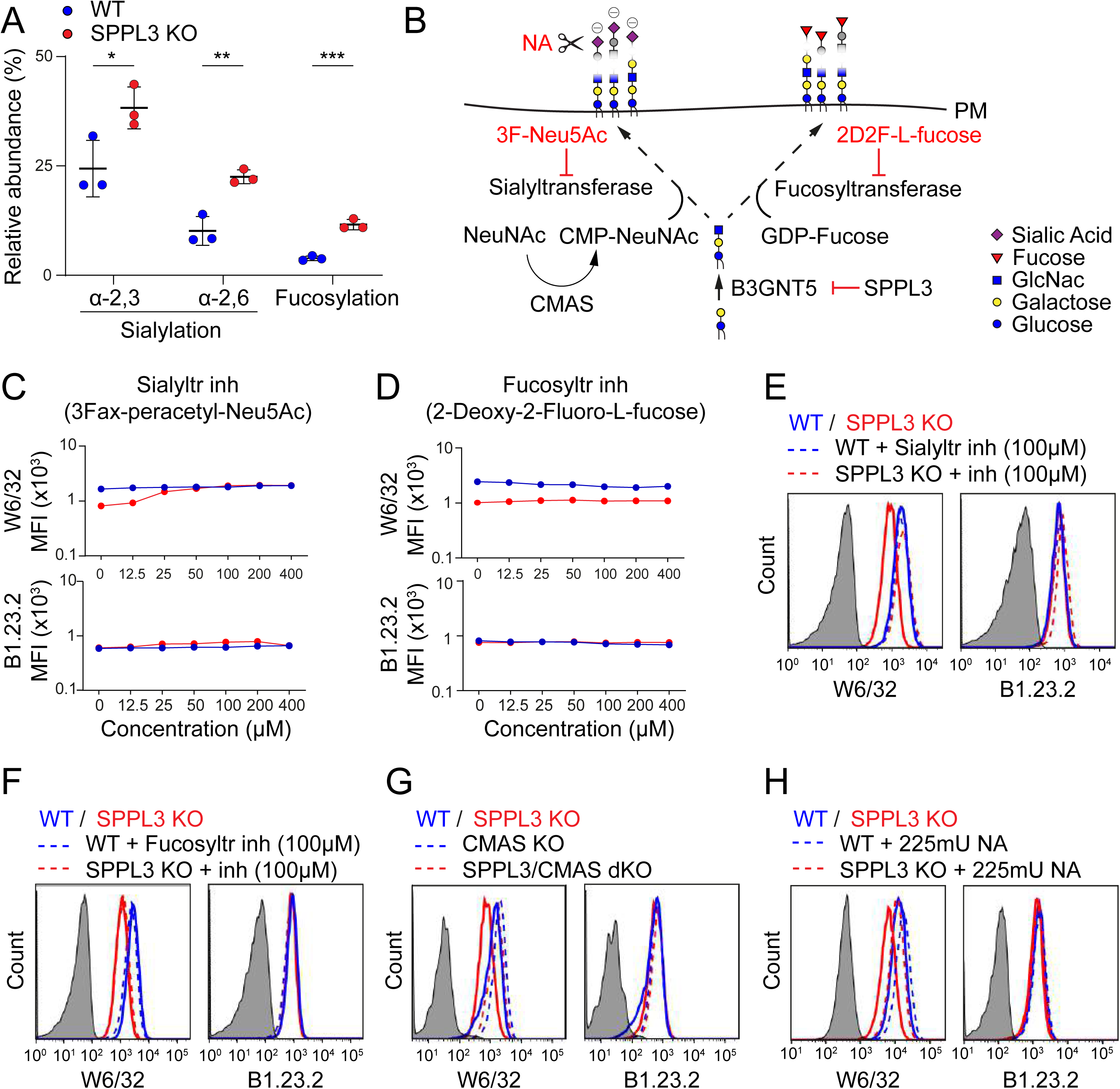
Sialic acid residues on nsGSLs are required for HLA-I shielding (A) Quantification of the percentage sialylated and fucosylated nsGSLs relative to total GSLs present in WT or SPPL3 KO HAP1 cells, n=3. Experiment described in Figure 5D. (B) Schematic model of several targetable steps in the sialylation and fucosylation of nsGSLs. NA, neuraminidase; PM, plasmamembrane. (C/D) Titrations of indicated sialyltransferease (C) and fucosyltransferase (D) inhibitors and their effect on W6/32 (*upper*) and B1.23.2 (*bottom*) cell surface staining of HAP1 WT (*blue*) and SPPL3 KO (*red*) cells. (E/F) Histograms of W6/32 (*left*) and B1.23.2 (*right*) cell surface staining of HAP1 WT (*blue*) or SPPL3 KO (*red*) cells with (*dashed*) or without (*continuous*) preincubation with indicated sialyl- (E) or fucosyltransferase (F) inhibitors (100 µM). Unstained control is in gray, representative for n=3. (G) Representative histograms of W6/32 and B1.23.2 cell surface stainings of HAP1 WT (*blue*), SPPL3 KO (red), CMAS KO (*blue dashed*) and SPPL3/CMAS double KO (dKO) cells (*red dashed*). CMAS KO cells were puromycin selected resulting in a pooled population of CMAS KO cells. Unstained control is in gray, n=2. (H) Representative histograms of W6/32 and B1.23.2 cell surface staining of HAP1 WT (*blue*) and SPPL3 KO (*red*) treated with (*dashed*) or without (*solid*) 225mU Neuraminidase (NA) for 1h at 37 degrees. Unstained control is in gray, n=3.

### Pharmacological inhibition of GSL synthesis in glioma enhances anti-tumor immune activation

Having determined that nsGSL-rich target cells suppress T cell activity, we investigated whether tumors increase nsGSL expression or downmodulate SPPL3 activity to escape T cell immunity. Because of the complexity inherent to identifying (large) nsGSLs, there is currently only a limited amount of data available on their tissue expression, including tumors (Merrill and Sullards, 2017; Zhang et al., 2019). Nonetheless, elevated levels of nsGSLs or its synthesis enzyme B3GNT5 have been observed on several tumor types including glioma, AML and adenocarcinomas (Furukawa et al., 2015; Hakomori, 1984; Wang et al., 2012; Wikstrand et al., 1991). In addition, The Cancer Genome Atlas (TCGA) analyses demonstrated that high B3GNT5 expression in low grade glioma correlates with decreased overall patient survival (Figure 7A). In line with our findings, the reverse holds true for the B3GNT5-suppressing SPPL3 (Figures 5B and 7B). Moreover, analyses of the combined effect of B3GNT5 and SPPL3 expression showed only lower survival rates for patients with high B3GNT5 and low SPPL3 expression (Figures 7C and S7A), probably reflecting that the nsGSL levels are only elevated in tumors from this group. This indicates that gliomas possibly escape immunity by exploiting the SPPL3-B3GNT5 axis. To demonstrate a role for this novel pathway in glioma immune evasion, we first assessed whether the function of SPPL3 to maintain HLA-I accessibility is lacking. We found that overexpression of SPPL3 in the glioblastoma cell line U373 induced an increase in HLA-I accessibility without altering HLA-I expression (Figure 7D). Next, we analyzed the role of GSLs in this process by genetic depletion of (ns)GSLs from WT U373 cells and detected also here a specific increase in HLA-I accessibility (Figure 7E). Moreover, in the absence of GSLs, U373 cells increased their capacity to engage the HLA-I ligand LIR-1 and were better activators of T cells (Figures 7F and 7G).

**Figure 7.**
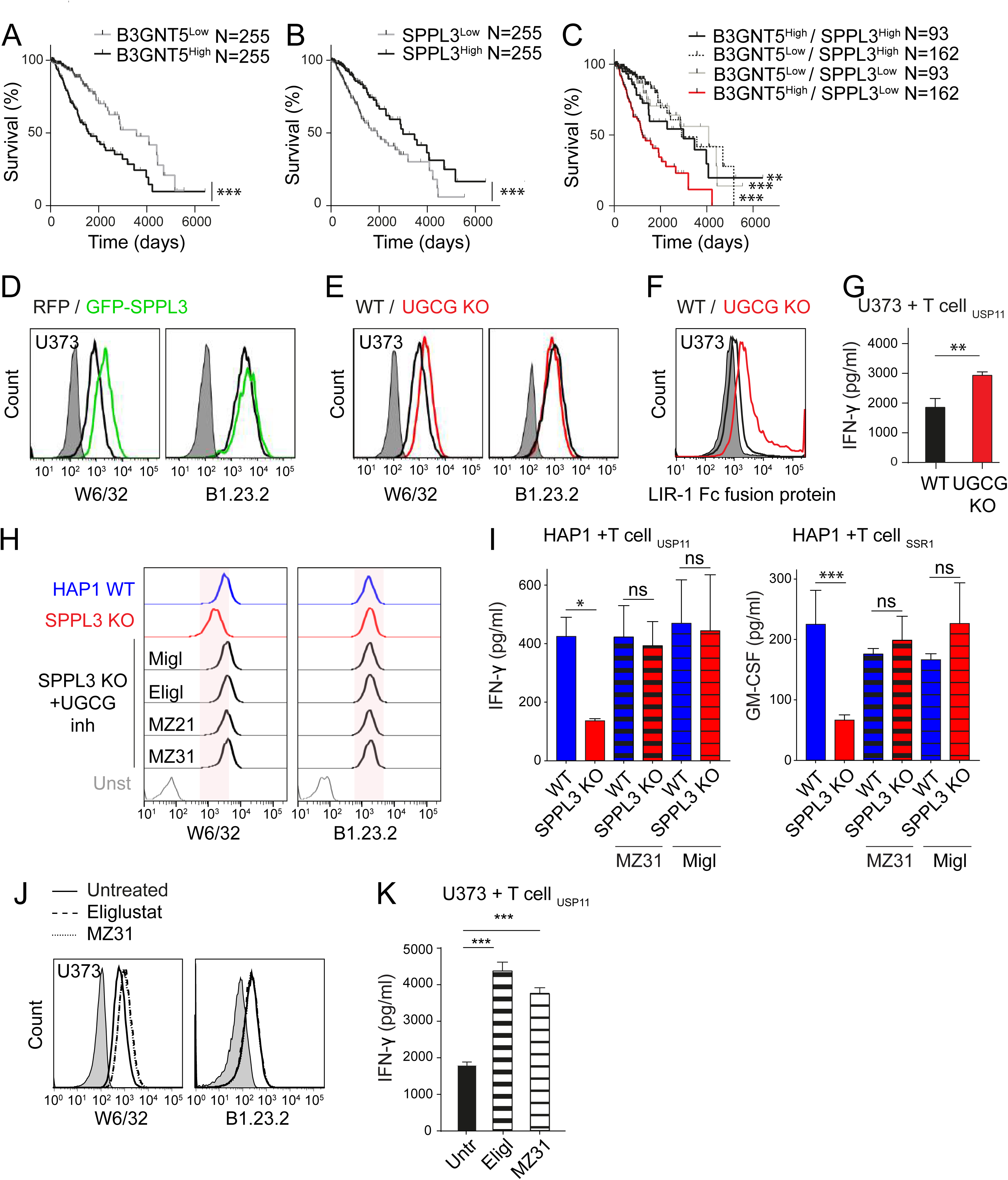
Pharmacological inhibition of GSL synthesis in glioma enhances anti-tumor immune responses. (A/B) TCGA (The Cancer Genome Atlas) derived Kaplan-Meier curve showing the percentage survival of patients have tumors expressing high (*black*) or low (*gray*) levels of B3GNT5 (A) or SPPL3 (B). (C) Kaplan-Meier curves for the four combined B3GNT5/SPPL3 expression levels (see Figure S7A for expression distribution). (D) Representative histograms of W6/32 (*left*) and B1.23.2 (*right*) cell surface staining of U373 glioblastoma cells overexpressing GFP-SPPL3 (*green*) or RFP empty vector control (*black*), mixed and analyzed in a single well. Unstained control in gray, n=2. (E) Representative histograms of W6/32 and B1.23.2 cell surface staining of WT (*black*) and UGCG KO (*red*) U373 cells. UGCG KO cells were puromycin selected resulting in a pooled population of KO cells. Unstained control is in gray, n=3. (F) Representative histogram of LIR-1 Fc fusion protein binding to U373 WT (*black*) and UGCG KO cells (*red*). Unstained control is in gray. n=2 (G) IFN-γ secretion by HLA-A*0201-restricted T cells recognizing USP11 peptides after overnight co-culture with U373 WT (*black*) or UGCG KO (*red*) cells as determined by ELISA. Representative plot of triplicate cultures are shown, n=3. (H) Representative histograms of W6/32 (*left*) and B1.23.2 (*right*) cell surface staining of HAP1 WT (*blue*), SPPL3 KO (*red*) and SPPL3 KO cells preincubated with UGCG inhibitors miglustat (Migl), eliglustat (Eligl), MZ21 or MZ31 (*black*). Unstained control is in gray. The red area indicates fluorescence of SPPL3 KO cells. n=3 (I) IFN-γ (left) or GM-CSF (right) secretion by HLA-A*0201-restricted T cells specific for endogenously derived USP11 (n=3), or SSR1 (n=1) antigens in an overnight co-culture with HAP1 WT (*blue*) or SPPL3 KO cells (*red*) each pre- incubated with (*dashed*) or without (*solid*) the indicated UGCG inhibitor (MZ31 or miglustat (Migl)) as determined by ELISA. Representative plots of triplicate cultures are shown. (J) Representative histograms of W6/32 (*left*) and B1.23.2 (*right*) cell surface staining of WT (*black, solid*) U373 cells preincubated with UGCG inhibitors eliglustat (*black, dashed*) or MZ31 (*black, dotted*). Unstained control is in gray, n=3. (K) IFN-γ secretion by HLA-A*0201-restricted T cells recognizing USP11 peptides after overnight co-culture with untreated (*black*) or depicted UGCG inhibitor pretreated (*dashed*) U373 cells as determined by ELISA. Representative plots are shown, n=2.

To regulate nsGSL dependent HLA-I driven immune responses in patients, the clinically approved GSL synthesis inhibitors miglustat and eliglustat may be used (Stirnemann et al., 2017). These drugs are currently being used as substrate reduction therapy in Gaucher disease. We first explored whether these small molecule drugs affect accessibility of HLA-I epitopes that are shielded in SPPL3 KO cells. The miglustat mimics MZ21 and MZ31 with fewer off-target effects were also included (Ghisaidoobe et al., 2014). All GSL synthesis inhibitors fully restored HLA-I accessibility despite a small proportion of GSLs still being detectable on the cell surface (Figures 7H, S7C and S7D). Moreover, these inhibitors increased the capacity of SPPL3 KO cells to activate T cells (Figures 7I and S7E). The same was true for the U373 cells of which HLA-I shielding was alleviated and their capacity to activate T cells was increased (Figures 7J, 7K and S7F). Together these data demonstrate the potential use of these inhibitors as a means to boost immune responses against tumors that display an excess of nsGSLs.

## DISCUSSION

The process of HLA-I antigen presentation has been a topic of longstanding interest to the research community, giving rise to a detailed understanding of various proteins governing this complex pathway. We here add an unexpected element to the equation of successful antigen presentation, namely the SPPL3-B3GNT5 pathway responsible for the production of a subset of GSLs. GSLs are present on every cell and yet their functional roles in the cell membrane remain largely unknown. By conducting sensitive genome-wide screens in an iterative fashion, we here uncovered a novel role for a subset of GSLs in immunity controlled by the protease SPPL3. These so-termed nsGSLs shield HLA-I molecules, causing diminished receptor-ligand interactions and decreased CD8^+^ T cell responses. We identified the aspartyl protease SPPL3 as a new switch controlling the expression of nsGSLs through proteolytic inhibition of the nsGSL synthesizing enzyme B3GNT5. Taken together, our study reveals a layer of immune regulation which acts in all likelihood independently of previously known components of the HLA-I antigen presentation machinery.

Various malignant cells exhibit alterations in their GSL surface repertoire to which a number of specific functions have been attributed. Some GSLs can serve as signaling molecules to control cellular processes such as apoptosis and proliferation while other GSL species can confer anti-cancer drug resistance by inhibiting proteins that facilitate their membrane transport (Liu et al., 2013; Ogretmen and Hannun, 2004). We here introduce the concept that changes in the tumor GSL repertoire, in particular increments of sialic acid containing nsGSLs, prevent HLA-I from signaling to T cells as means to evade immune surveillance. Supporting this theory, CD8^+^ T cells show a GSL-mediated diminished capacity to respond to glioma cells, a tumor type with high levels of nsGSLs (Furukawa et al., 2015). Furthermore, survival analyses of glioma patients revealed the worst overall survival when the GSL synthesis signature of the tumor is skewed towards high nsGSL levels. Other tumor types, including AML, colorectal carcinoma, adenocarcinomas and ductal carcinomas in situ (DCIS) also overexpress B3GNT5 and its product nsGSLs, suggesting that HLA-I shielding is a more general strategy for tumor immune evasion (Hakomori, 1984; Potapenko et al., 2015; Wang et al., 2012; Wikstrand et al., 1991).

Also pathogens, such as cytomegalovirus, respiratory syncytial virus and HIV alter the GSL composition of the host cell, potentially inducing/representing immune evasion through HLA-I shielding (Fantini et al., 2000; Moore et al., 2008; Radsak and Wiegandt, 1984). Except for low resolution data of cytomegalovirus-induced nsGSL expression upon infection (Andrews et al., 1989; Radsak and Wiegandt, 1984), currently little is known about infection-induced complex nsGSL expression.

Understanding nsGSL function at a molecular level and in (patho)physiological settings is challenging given that their isolation, analytical dissection and in particular their experimental manipulation are extraordinarily demanding to date (Merrill and Sullards, 2017; Zhang et al., 2019). Hence, no validated methods are available to study nsGSL-protein interactions restricting options to study the nsGSL-HLA-I interaction. Still, our data indicate that the interaction between nsGSLs and HLA-I molecules must be transient since a high dose of antibody can overcome the decreased accessibility. In addition we show that carbohydrate-carbohydrate interactions (D’Angelo et al., 2013) between nsGSLs and HLA-I *N*-glycans are unlikely to play a role in HLA-I shielding (Figure S2). Profound shielding of large HLA-I areas by nsGSLs, as we describe here, can however be explained by the fact that, in contrast to other GSL subtypes, nsGSLs can carry huge glycan chains of up to 60 sugar residues (Miller-Podraza et al., 1993; Miller-Podraza et al., 1997). These long chains may reach up to HLA-I domains needed for interaction with ligands such as LIR-1. The nsGSL-HLA-I interaction may cause steric hindrance for molecular interactions with HLA-I, mediated by the nsGSLs themselves, or by altered HLA-I orientation towards the cellular membrane (Mitra et al., 2004). In addition, we show that sialic acid residues on GSLs are essential for HLA-I shielding. The negatively charged sialylated nsGSLs may establish an ionic interaction with HLA-I, which has abundant positively charged patches at its molecular surface (Li et al., 2012). Similar interactions have been found between sialic acids on small GSLs and exposed positively charged amino acid residues right above the plasma membrane (D’Angelo et al., 2013). Finally, our data do not exclude a direct interaction between the GSL ceramide and the HLA-I transmembrane domain (Contreras et al., 2012) which could contribute to positioning the nsGSL glycan chain in close proximity of HLA-I.

In this study, we present GSLs as highly relevant molecules affecting the efficiency of immune responses. nsGSLs and its molecular switch SPPL3 therefore represent an unexplored avenue for therapeutic intervention in pathological conditions where HLA-I antigen presentation plays a central role, such as cancer, infection and autoimmune diseases. Currently, two small molecule drugs inhibiting GSL synthesis are registered, miglustat (Zavesca) and eliglustat (Cerdelga). These structurally different UGCG inhibitors (Platt et al., 1994; Shayman, 2010) have been approved for the treatment of patients with lysosomal storage disorders, such as type I Gaucher disease and Niemann- Pick disease type C (Lachmann, 2003; Wraith and Imrie, 2009). Therapeutic application can therefore efficiently be extended to include immune enhancement against tumors or pathogen-infected cells through improved HLA-I accessibility. GSL synthesis inhibition may even be successfully combined with existing in human immunotherapies, such as PD-1 blockade, because of the potential synergy between enhanced tumor cell immunogenicity and simultaneous T cell activation. Hence our findings define a novel strategy to improve immunotherapy.

## MATERIALS AND METHODS

Descriptions of used cell lines, culture conditions, antibodies, recombinant proteins, inhibitors, gRNA sequences, DNA constructs, flow cytometry, immunoprecipitation, SDS-PAGE, western blotting, Sanger sequencing, siRNA transfection, viral production/transduction, in-gel tryptic digestion, LC-ESI-MS/MS of glycopeptides, B3GNT5 activity assay,crystal structures and TCGA analyses can be found in the *Extended Materials and Methods section*.

### Haploid genetic screening

Genome-wide knockout screening was performed in either early passage WT or CRISPR/Cas9 generated SPPL3 KO HAP1 cells using directly conjugated W6/32 or BB7.2 antibodies. Retroviral mutagenesis was performed on ∼100 x 10^6^ cells using GT-GFP or GT- BFP plasmids as previously described (Brockmann et al., 2017). Around 2 x 10^9^ expanded mutagenized cells were fixed, stained and sorted (Brockmann et al., 2017). Integration sites were amplified using LAM-PCR on genomic DNA isolated from sorted cells and analyzed by a deep sequencing approach as described (Brockmann et al., 2017). An extended description can be found in the *Extended Materials and Methods section*.

### Genome editing

SPPL3 KO and tapasin KO HAP1 cells were created by in frame integration of a blasticidin-resistance gene after cotransfection of pX330 with TIA-2Ablast (using Extremegene HP, Sigma) as described for other targets (Blomen et al., 2015). Lentiviral constructs containing gRNAs targeting UGCG, the five core GSL-enzymes, CMAS and GANAB were co-transfected into HEK293T with the packaging enzymes psPAX2, pVSVg, pAdVAntage using polyethylenimine (PEI; Polyscience) for virus production. Filtered viral supernatants were used for transduction by spinoculation in the presence of 8µg/mL protamine sulfate. Cells were selected using puromycin (0.25µg/mL; Gibco), blasticidin (10µg/mL; Gibco), or gated based on the co-expression of GFP. KO clones for SPPL3, tapasin, UGCG and GANAB were made by limiting dilution and sequence verified. HLA-A, -B and -C KO cells were generated by pX330 transfection followed by single cell FACS sort using W6/32 and sequence verification of clones (de Waard, 2020). Additionals details can be found in the *Extended Materials and Methods section*.

### T cell assays

Target cells were co-cultured with T cells in a 1:1 ratio for 18h as previously described (Spaapen et al., 2008). Cytokine content of cell free supernatants was determined using standard sandwich ELISA according to the manufacturer’s instructions (IFN-γ and GM-CSF; Sanquin and Biolegend). Some target cells were pre-incubated for 2 days in the presence or absence of UGCG inhibitors.

### GSL LC-MS

GSLs were extracted from 1 x 10^7^ HAP1 WT, SPPL3 KO or SPPL3/B3GNT5 double KO cells and purified by RP-SPE. GSL glycans were released by EGCase I and purified. Reduction and GSL glycan desalting was carried out as described previously with slight modifications (Jensen et al., 2012). After carbon SPE clean-up, the purified glycan alditols were analyzed by porous graphitized carbon (PGC) LC-ESI-MS/MS. GSL-derived glycan structures were assigned based on the MS/MS spectra, SphingoMAP and specific software tools. Detailed descriptions can be found in the *Extended Materials and Methods section*.

### Statistical analyses

All error bars correspond to the standard deviation of the mean. Data from genome-wide screens were analyzed using two-sided Fisher’s exact test followed by FDR (Benjamini-Hochberg) correction of the p-value. Other statistical evaluations were done by a Student’s t-test (analysis of two data groups), one-way ANOVA (three groups or more), two-way ANOVA (two variables), Mann-Whitney U test (non-parametric analyses) or Log-rank (survival analyses) with Prism software (http://www.graphpad.com). *p<0.05, **p<0.01, ***p<0.001 and ns (= not significant). EC50 values of titrations were calculated using non-linear four parameter fit modeling with Prism software.

## ACKNOWLEDGMENTS

We thank Dr. R. Fluhrer (Ludwig Maximilians University, Germany) for providing SPPL3 plasmids, Dr. M. Griffioen (LUMC, The Netherlands) for providing T cell clones, Dr. O. Mandelboim (Hebrew University Hadassah Medical School, Israel) for providing LIR-1 Fc fusion protein, Dr. T. Rispens (Sanquin, The Netherlands) for expert advice and assistance on protein purification and Fab generation, E. Mul, S. Tol and M. Hoogenboezem (Sanquin) for assistance with flow cytometry, Dr. I. Berlin (LUMC) for critical reading of the manuscript, Dr. Y. Rombouts (Université de Toulouse, France) for discussions and contribution to MTBE extraction, and C.A.M. Koeleman and A.L. Hipgrave Ederveen (LUMC) for technical support with LC-MS.

This work was supported by the Netherlands organization for scientific research (NWO-VENI 016.131.047 and ZonMw-ETH 435004024; R.M.S.), KWF Alpe d’HuZes (Bas Mulder Award 2015- 7982; R.M.S.) and the Landsteiner Foundation for Blood Transfusion Research (LSBR fellowship 1842F; R.M.S.). M.L.M.J. was supported by an EMBO short term fellowship (ASTF 10-2016). T.R.B. was supported by NWO Vici Grant 016.Vici.170.033, KWF grant NKI2015-7609, the Cancer Genomics Center (CGC.nl), and the Ammodo KNAW Award 2015 for Biomedical Sciences. T.R.B. is a co-founder and advisory board member of Haplogen GmbH and co-founder and managing director of Scenic Biotech B.V.. S.F. was supported by grants from DOD (W81XWMH-16-1-0500) and NIH (R01DE028172, R03CA216114, RO3CA223886 and RO3CA231766). The authors have no conflicting financial interests. Processed screen results are accessible in an interactive database (https://phenosaurus.nki.nl/).

## AUTHOR CONTRIBUTION STATEMENTS

Conceptualization and design, M.L.M.J and R.M.S. Data acquisition, analysis and interpretation, M.L.M.J, M.R., A.A.W., T.Z., B.C., R.P., V.A.B., A.X., T.V., S.B., L.J., E.S., S.H., R.P. and R.M.S. Resources and discussion, A.M., S.F., F.H.J.C., M.H.M.H., M.G. and H.O. Supervision and conceptual discussion, J.B.H., M.W., T.R.B., J.N., R.M.S. Writing, M.L.M.J. and R.M.S. Editing, J.B.H., M.R., A.A.W.,T.Z., M.W., J.N.

---

**Table S1.**
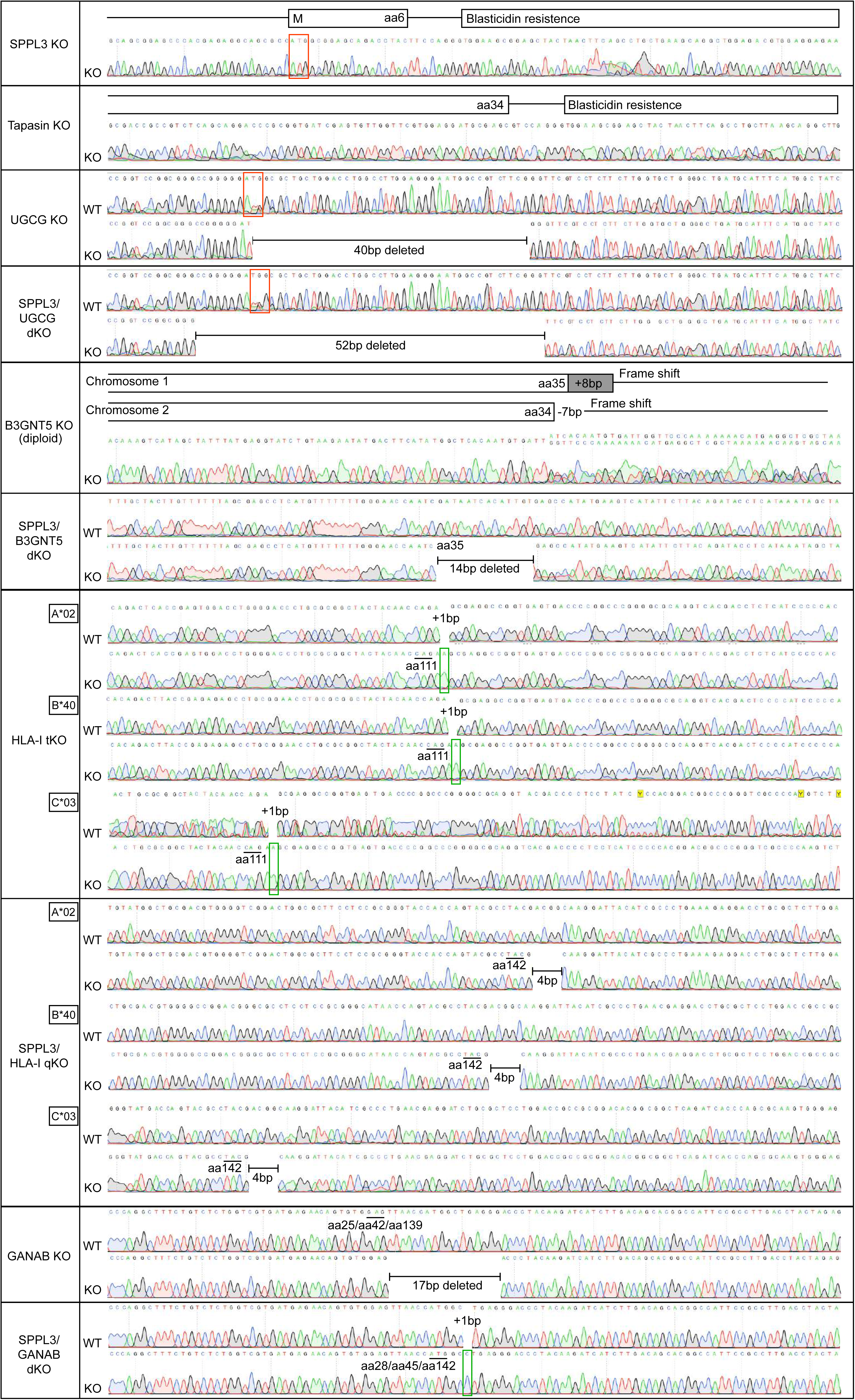
**Sequences of CRISPR/Cas9 generated monoclonally derived KO cell lines (Relates to Figures 1, 2, 3, 4, 5 and 6)** Genomic DNA isolated from the indicated monoclonally derived KO cell lines was amplified by PCR of the gRNA targeted region and Sanger sequenced.

**Table S2.**
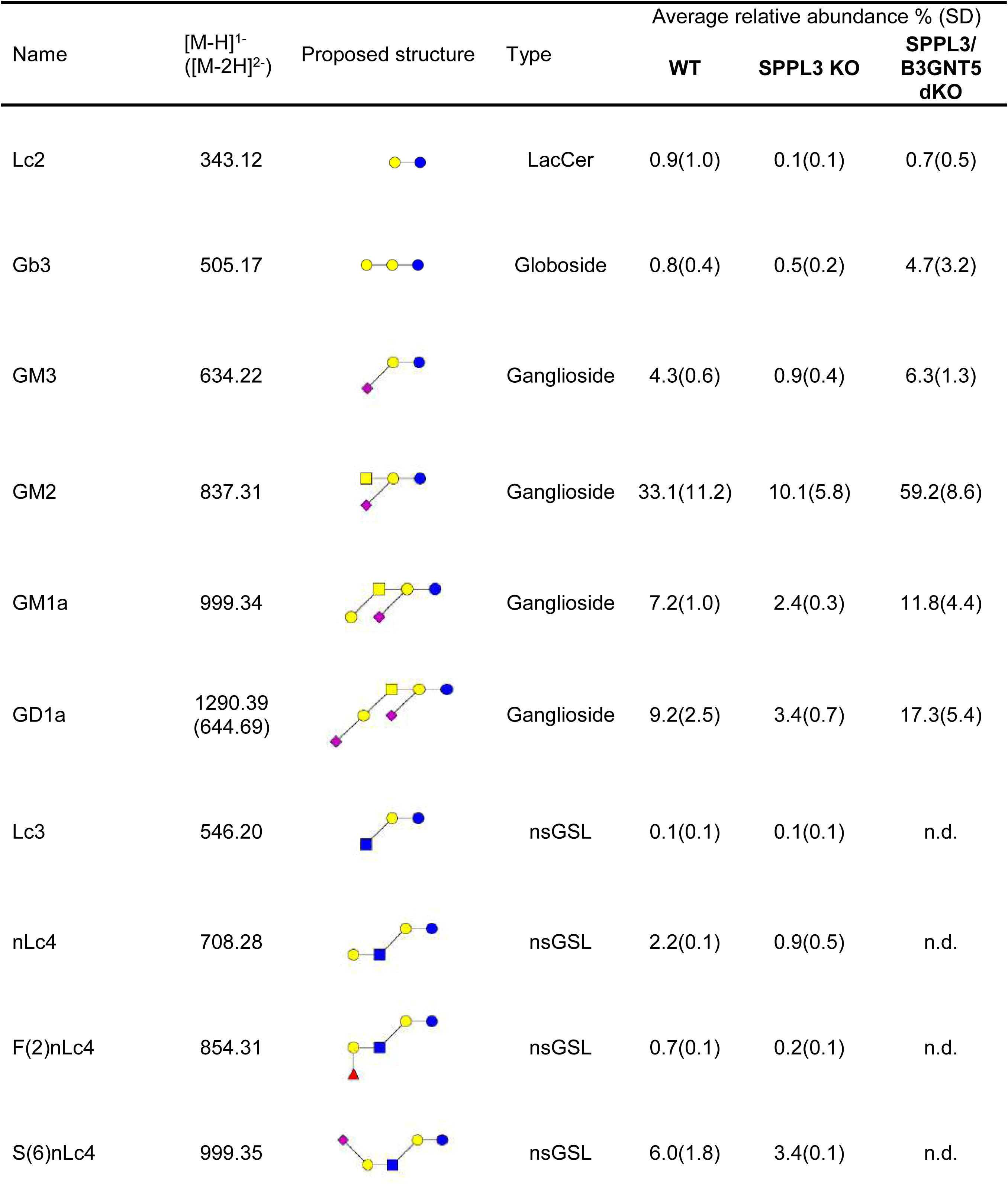
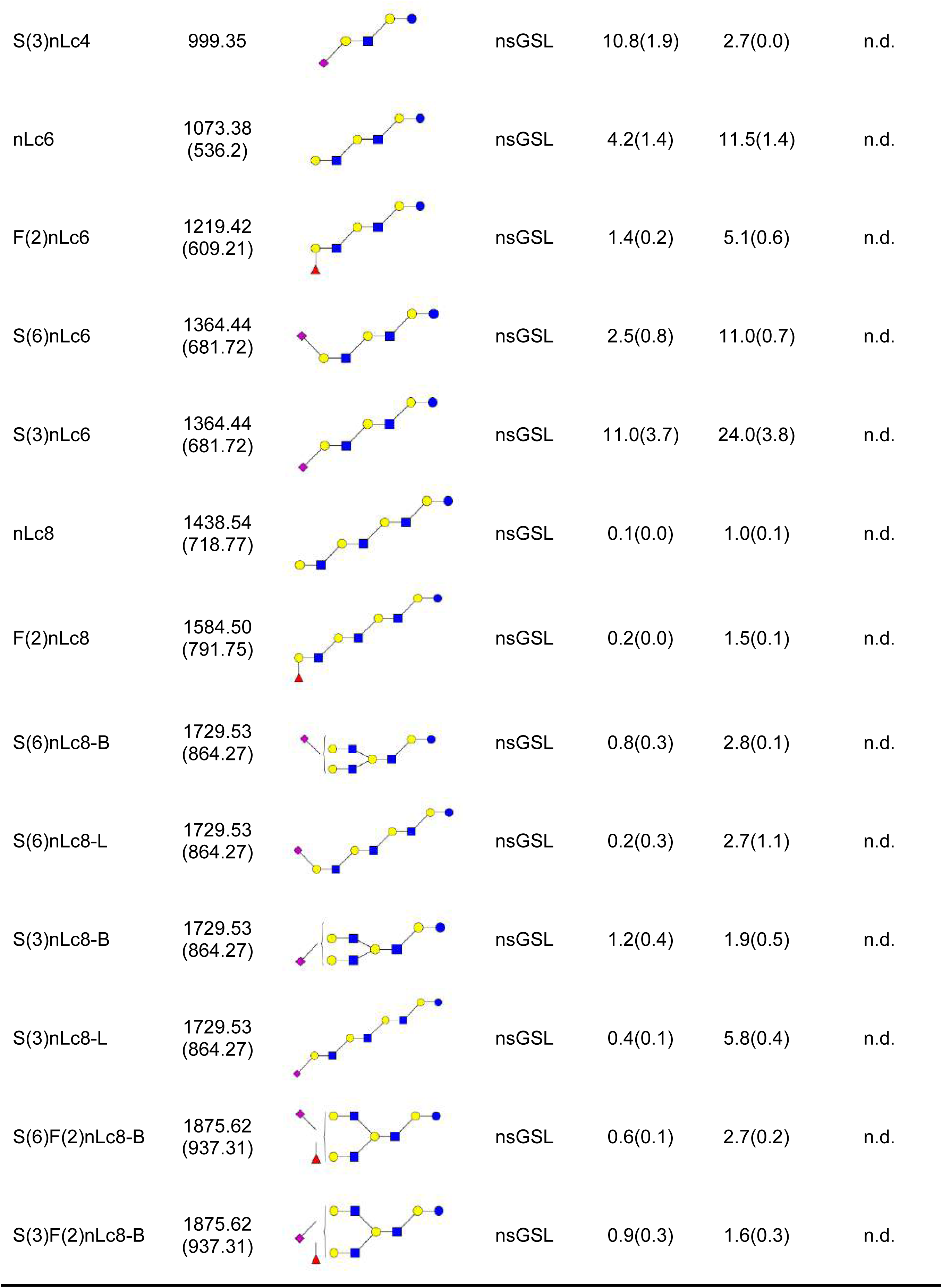
**Relative quantification of glycans released from GSLs of WT, SPPL3 KO and SPPL3/B3GNT5 double KO (dKO) cells using PGC LC-ESI-MS/MS** (Relates to Figure 5) Proposed structures were assigned based on MS/MS fragmentation (where possible) and biological GSL pathway constraints. Structures are depicted according to the CFG (Consortium of Functional Glycomics). Average from n=3, not detected (n.d.)

**Table S3.**
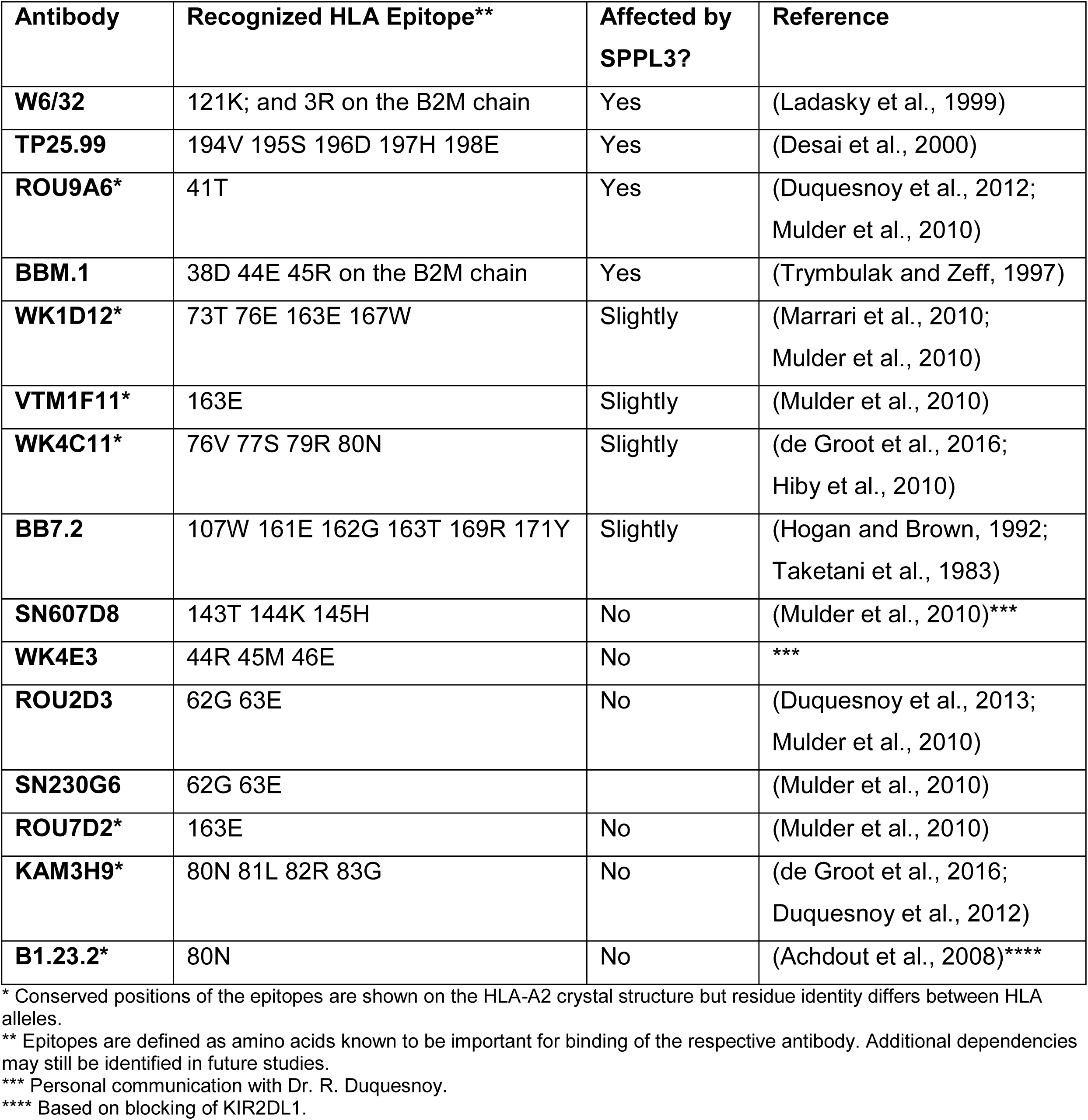
Epitopes of HLA-I specific antibodies

**Table S4.**
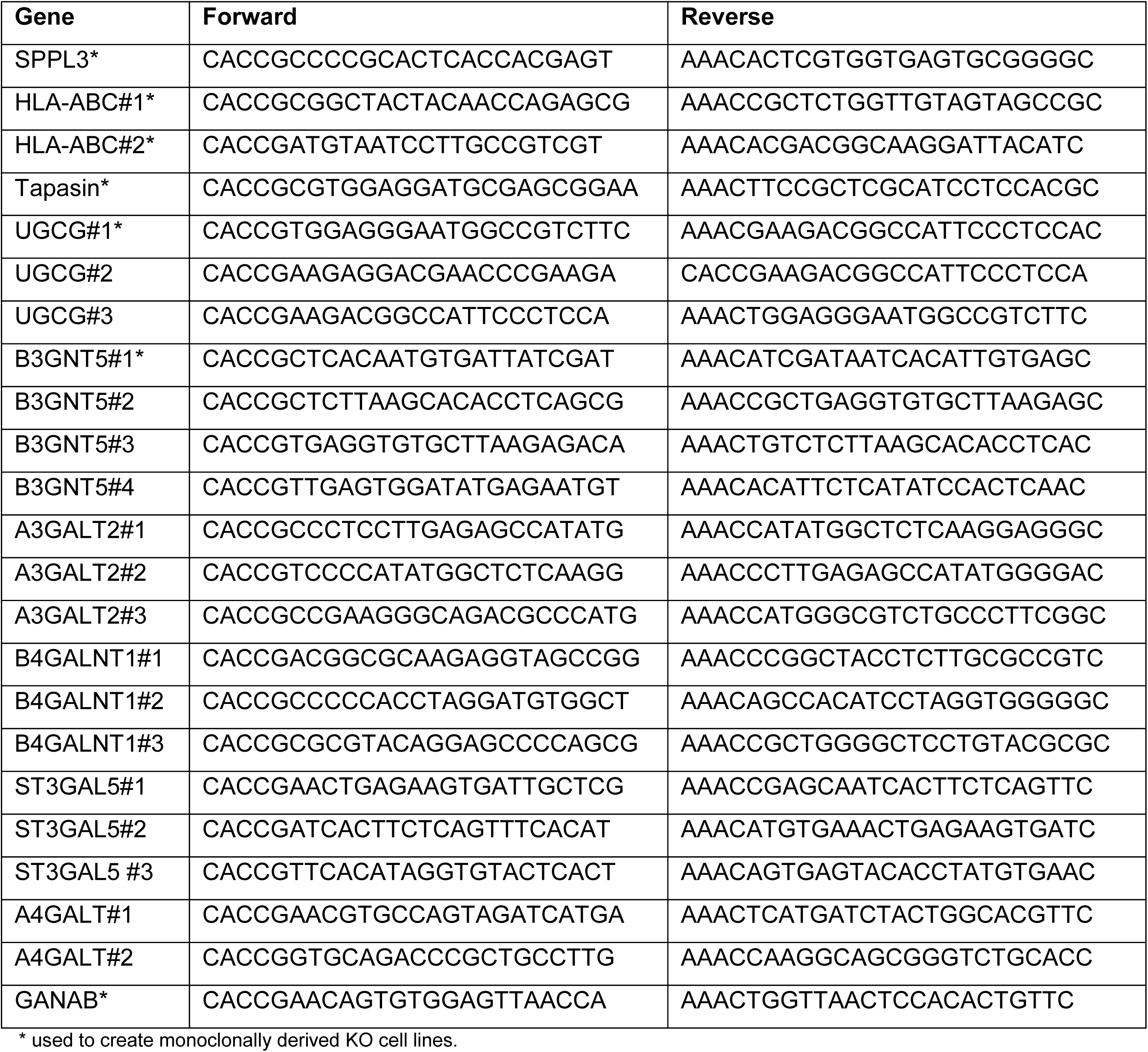
**gRNA sequences used for CRISPR/Cas9 mediated genome editing** gRNAs were handpicked using online tools (Doench et al., 2016; Hsu et al., 2013). The HLA-A, -B, -C specific gRNAs target common sequences of the three HLA alleles expressed in HAP1 cells (HLA-A*02:01, HLA-B*40:01 and HLA-Cw*03:04).

**Table S5.**
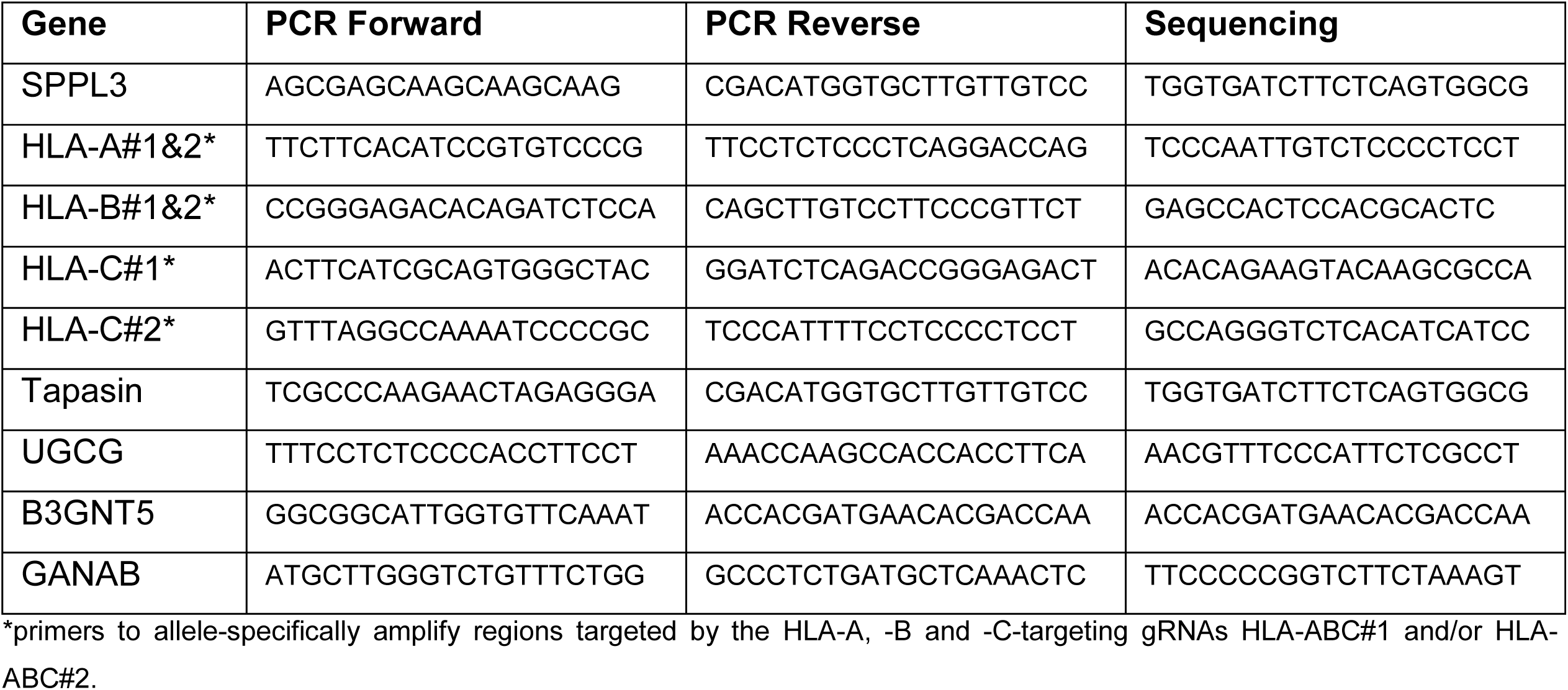
PCR and sequencing primers

**Figure S1.**
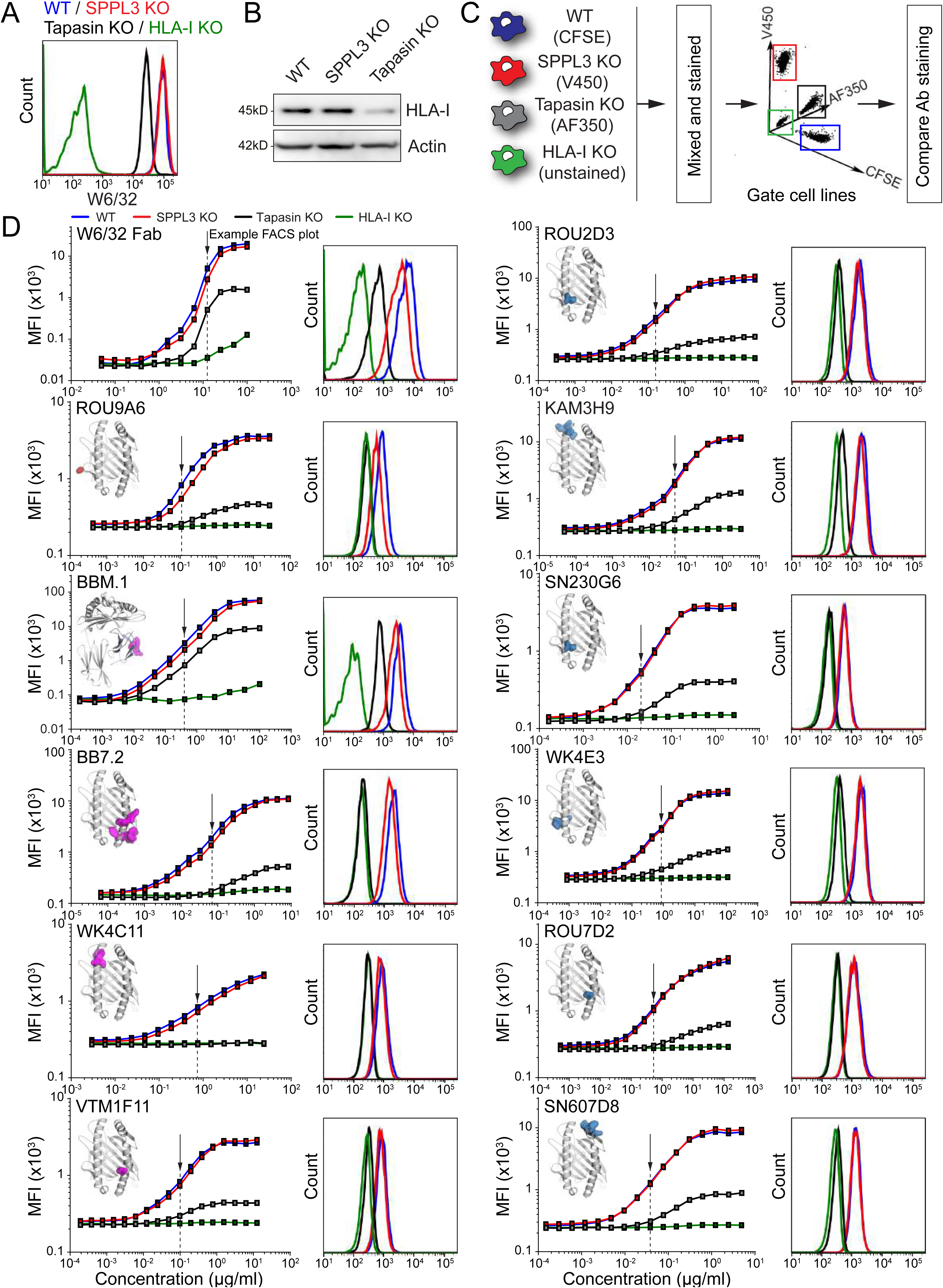
**SPPL3 activity determines the accessibility of membrane proximal HLA-I regions (*Relates to*** Figure 1) (A) Representative histogram of saturating W6/32 stain of HAP1 WT (*blue*), SPPL3 KO (*red*), tapasin KO (*black*) and HLA-I KO cells (*gray filled*). n=3 (B) Immunoblot showing the levels of total HLA-I (HC10) and actin (AC-15, loading control) in WT HAP1, SPPL3 KO and tapasin KO cells. n=2 (C) Schematic overview of fluorescent cell barcoding procedure. The depicted cell lines were barcoded using Alexa Fluor 350, Violet or CFSE proliferation dyes, mixed and stained for flow cytometry. The cell lines were gated based on their proliferation dye before analysis of antibody staining. (D) (*left*) Representative titration curves of AF647-labeled W6/32 Fab fragments and 11 HLA-I / B2M-specific antibodies on HAP1 WT (*blue*), SPPL3 KO (*red*), tapasin KO (*black*) and HLA-I KO (*green*) cells. Each inset highlights the epitope recognized by the respective antibody mapped on the HLA-I / B2M crystal structure in red (SPPL3-susceptible epitopes), purple (mildly affected epitopes) or blue (SPPL3-independent epitopes). (*right*) Representative histograms of non-saturating antibody staining (as indicated by the arrow) on HAP1 WT (*blue*), SPPL3 KO (*red*), tapasin KO (*black*) and HLA-I KO (*green*) cells are shown. n=3

**Figure S2.**
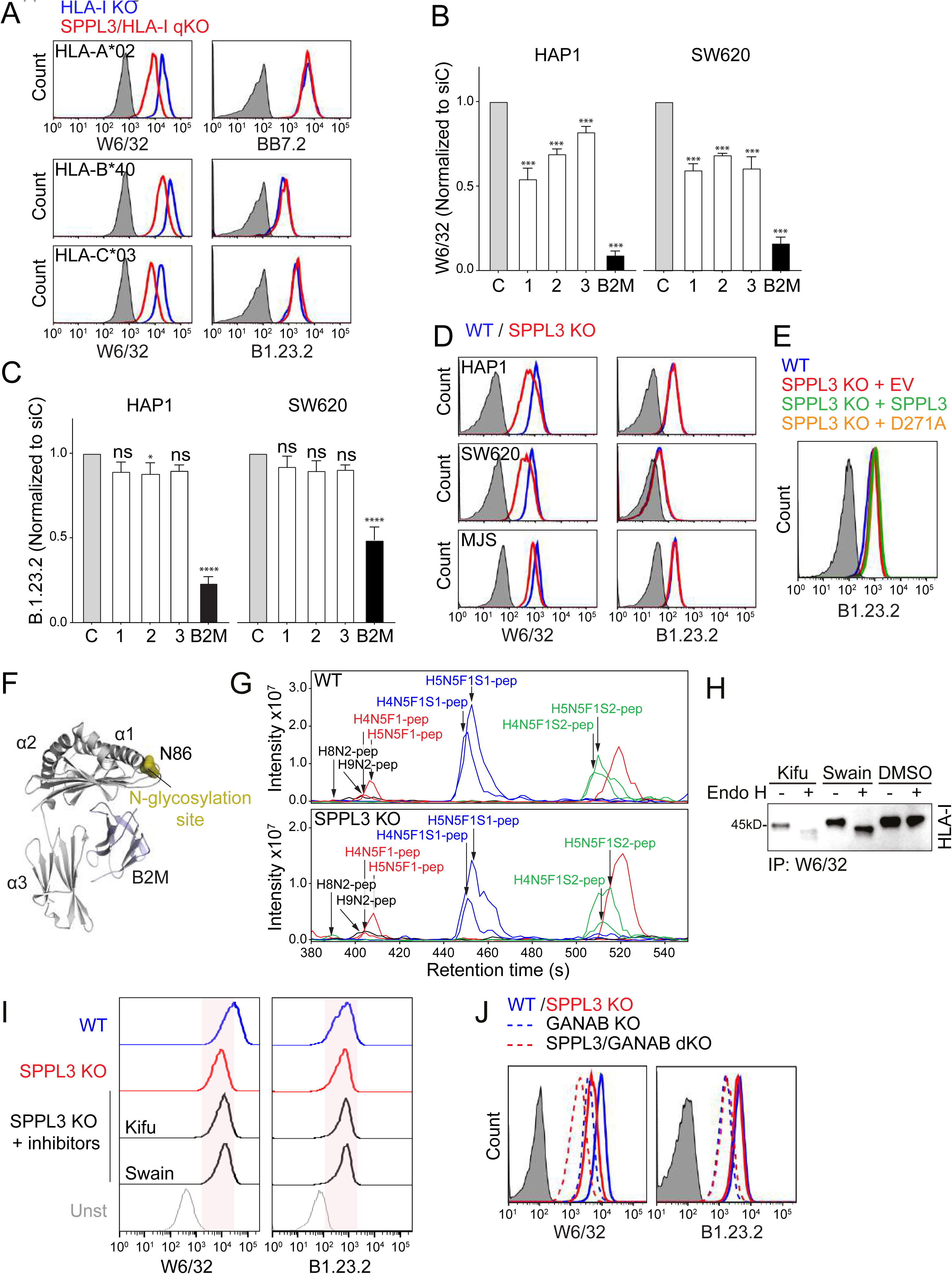
SPPL3 activity determines the accessibility of HLA-A, -B and -C alleles without affecting HLA-I N-glycosylation (*Relates to* Figure 2) (A) Representative histograms of W6/32 and BB7.2 (HLA-A) or B1.23.2 (HLA-B and -C) cell surface staining of HLA-A, -B and -C triple KO (HLA-I KO) (*blue*) or SPPL3/HLA-I quadruple KO (qKO) (*red*) HAP1 cells transduced with HLA-A*02:01, HLA-B*40:01 or HLA-C*03:03. n=1 (B/C) Mean fluorescent intensity of W6/32 (B) and B1.23.2 (C) cell surface staining of HAP1 or SW620 cells transfected with three individual siRNAs targeting SPPL3 or one siRNA targeting B2M relative to control (siC). n=2 (D) Representative histograms of W6/32 (*left panels*) and B1.23.2 stain (*right panels*) on either WT (*blue*) or SPPL3 gRNA (lentiCRISPR_v2) transduced and puromycin selected (*red*) HAP1, SW620 or MelJuSo cells. n=2 (E) Representative histogram of non-saturating B1.23.2 stain on HAP1 WT (*blue*) or SPPL3 KO cells transfected with either RFP-empty vector (*red*), RFP-SPPL3 (*green*) or catalytically inactive RFP-SPPL3 D271A (*orange*). n=2 (F) Crystal structure of HLA-I / B2M showing its N-glycosylation site. (G) Extracted ion chromatogram showing the 8 most abundant glycopeptides derived from tryptic digest of immunoprecipitated HLA-I (BB7.2) from HAP1 WT (*upper panel*) and SPPL3 KO (*bottom panel*) cells analyzed by nanoLC-ESI-IT-MS. The two most abundant high-mannose glycopeptides (*black*), neutral complex glycopeptides (*red*), monosialylated glycopeptides (*blue*) and disialylated glycopeptides (*green*) are shown. (H) Immunoblot showing HLA-I (HC10) immunoprecipitated by W6/32 from the depicted N-glycosylation inhibitor-treated SPPL3 KO cells followed or not by Endoglycosidase H treatment. (I) Representative histograms of W6/32 (*left*) and B1.23.2 (*right*) cell surface staining of HAP1 WT (*blue*), SPPL3 KO (*red*) and SPPL3 KO cells preincubated with N-glycosylation inhibitors kifunensine and swainsonine (*black*). n=3 (J) Histograms of W6/32 (*left*) and B1.23.2 stain (*right*) on either WT (*blue*), SPPL3 KO (*red*), GANAB KO (*blue dashed*) or SPPL3/GANAB double KO (dKO) cells (*red dashed*). Representative for n=3. For all flow cytometry data, the gray histogram represents an unstained control cell line.

**Figure S3.**
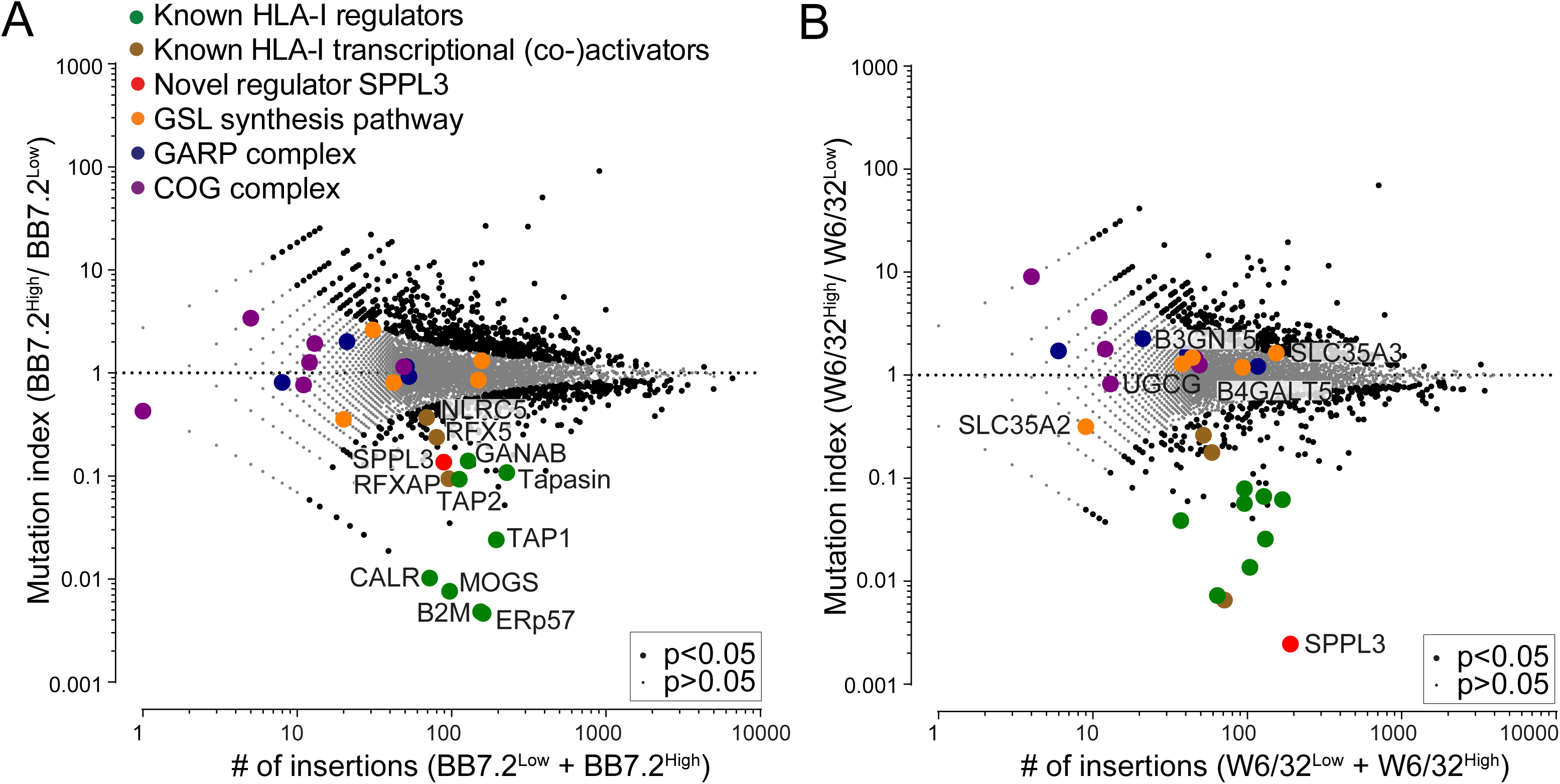
**Additional haploid genetic screens carried out using different HLA-I antibodies (*Relates to*** Figure 3) (A) Fish-tail plot: The mutation index shows the relative frequency of unique disrupting gene trap integrations in the BB7.2^High^ versus the BB7.2^Low^ sorted cell populations plotted against the total amount of unique disrupting integrations (BB7.2^High^ + BB7.2^Low^) mapped per gene. Positive and negative regulators of HLA-I (*black/color*) were determined by two-sided Fisher’s exact test, FDR (Benjamini-Hochberg) corrected p<0.05. Known HLA-I regulators are depicted in green, known HLA-I transcriptional activators in brown, the novel regulator SPPL3 in red, proteins involved in the glycosphingolipid synthesis pathway in orange and members of the GARP and COG complex in blue and purple respectively. (B) Fish-tail plot (same as in Figure 1B but with additional genes depicted): The mutation index shows the relative frequency of unique disrupting gene trap integrations in the W6/32^High^ versus the W6/32^Low^ sorted cell populations plotted against the total amount of unique disrupting integrations (W6/32^High^ + W6/32^Low^) mapped per gene. Positive and negative regulators of HLA-I (*black/color*) were determined by two-sided Fisher’s exact test, FDR (Benjamini-Hochberg) corrected p<0.05. Legend as in panel A.

**Figure S4.**
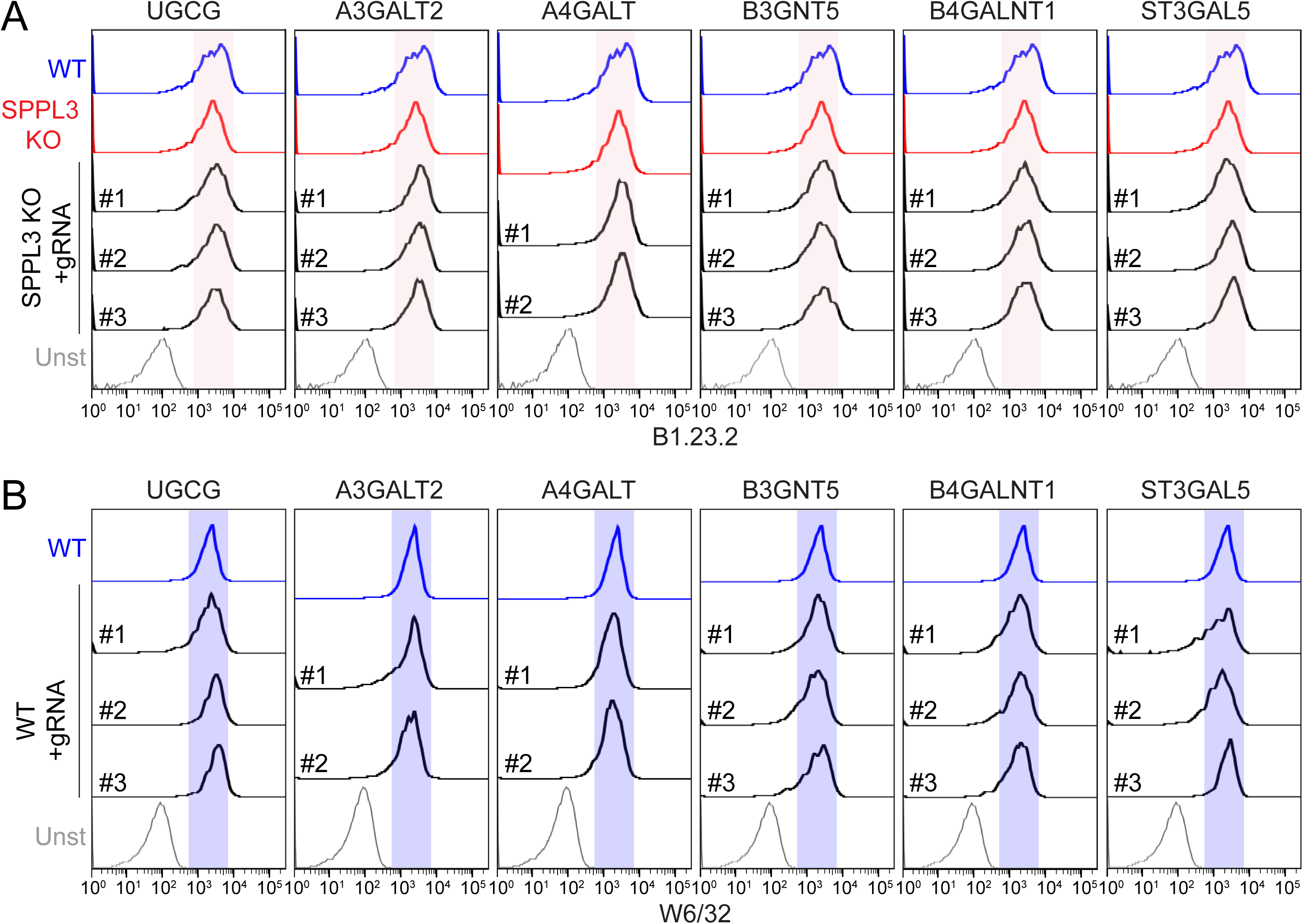
**HLA-I expression is not altered in cells lacking GSL synthesis enzymes (*Relates to*** Figure 4) (A) Representative histograms of non-saturating B1.23.2 cell surface stainings of HAP1 WT (*blue*), SPPL3 KO (*red*) or polyclonal populations of cells double KO (dKO) for SPPL3 and the core enzyme UGCG or the branching enzymes A3GALT2, A4GALT, B3GNT5, B4GALNT1 or ST3GAL5 (*black*). Double KO (dKO) cells were generated by CRISPR/Cas9 using two or three different gRNAs per gene in individual wells. Unstained control is in gray. Shown are transduced cells in the GFP^+^ gate. The red area indicates fluorescence of SPPL3 KO cells. n=2 (B) Representative histograms of non-saturating W6/32 cell surface staining of HAP1 WT (*blue*) or polyclonal populations of HAP1 cells KO for the core enzyme UGCG or the branching enzymes A3GALT2, A4GALT, B3GNT5, B4GALNT1 or ST3GAL5 (*black*). KO cells were generated by CRISPR/Cas9 using two or three different gRNAs per gene as indicated. Unstained control is in gray. Shown are transduced cells in the GFP^+^ gate. The blue area indicates fluorescence of WT cells. n=2

**Figure S5.**
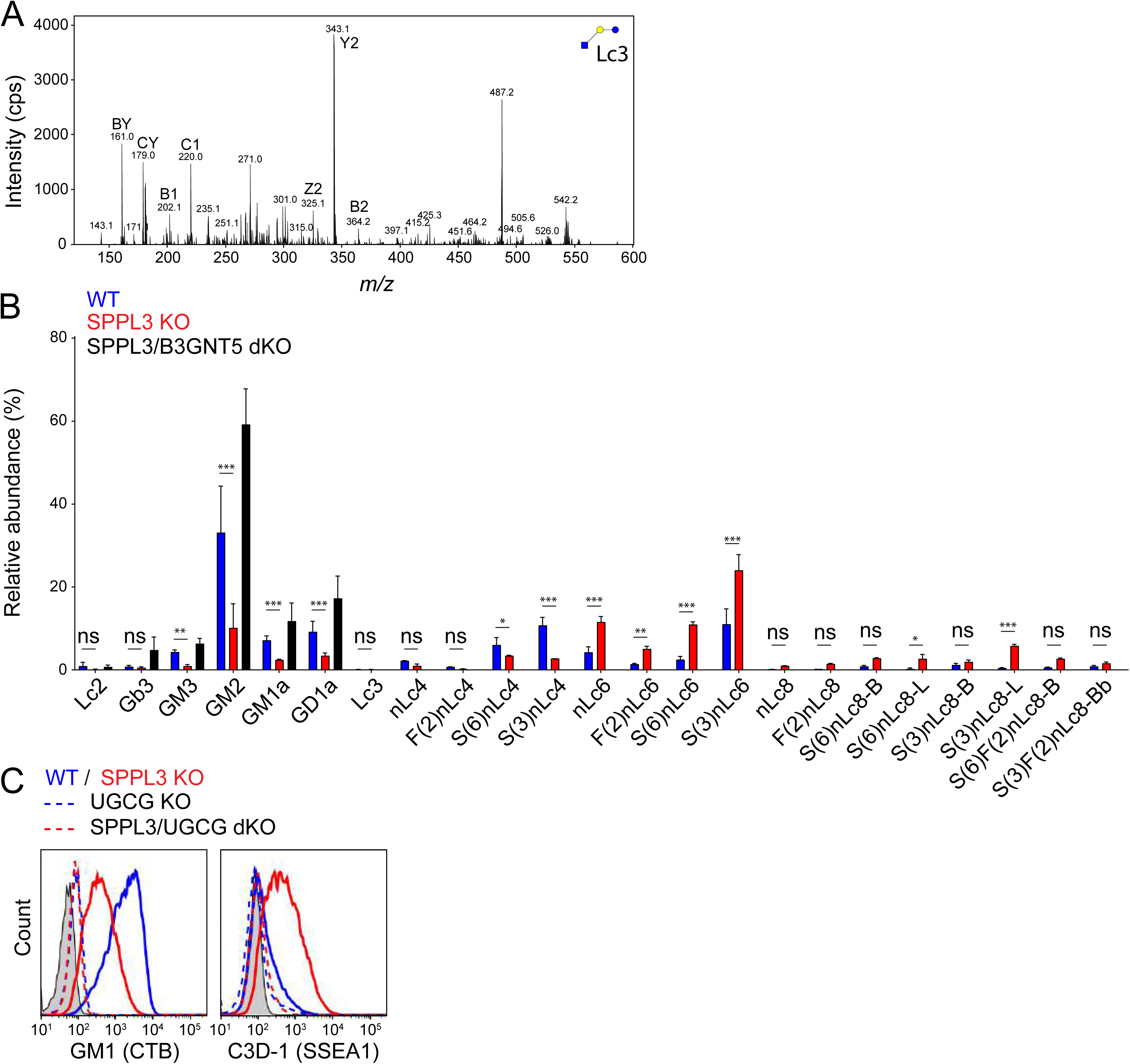
**Characterization of the GSL composition and effect on HLA-I accessibility of HAP1 cells (*Relates to*** Figure 5) (A) MS/MS spectra of the enzymatically released glycan from the lower band GSLs in the SPPL3 KO sample scratched from the TLC plate (Figure 5B), showing that this product is BODIPY-Lc3Cer. (B) Mean relative abundance of the 23 most abundant GSL glycans derived from WT, SPPL3 KO and SPPL3/B3GNT5 double KO (dKO) cells. Proposed structures were assigned based on MS/MS fragmentation (where possible) and biological GSL pathway constraints. Statistics shown compare GSL levels between WT and SPPL3 KO cells. n=3. (C) Representative histograms of flow cytometry of Cholera Toxin B (anti-GM1) and C3D-1 (SSEA-1) stained HAP1 WT (*blue*), SPPL3 KO (*red*), UGCG KO (*blue dashed*) and SPPL3/UGCG double KO (dKO) cells (*red dashed*). Unstained control is in gray. n=3

**Figure S6.**
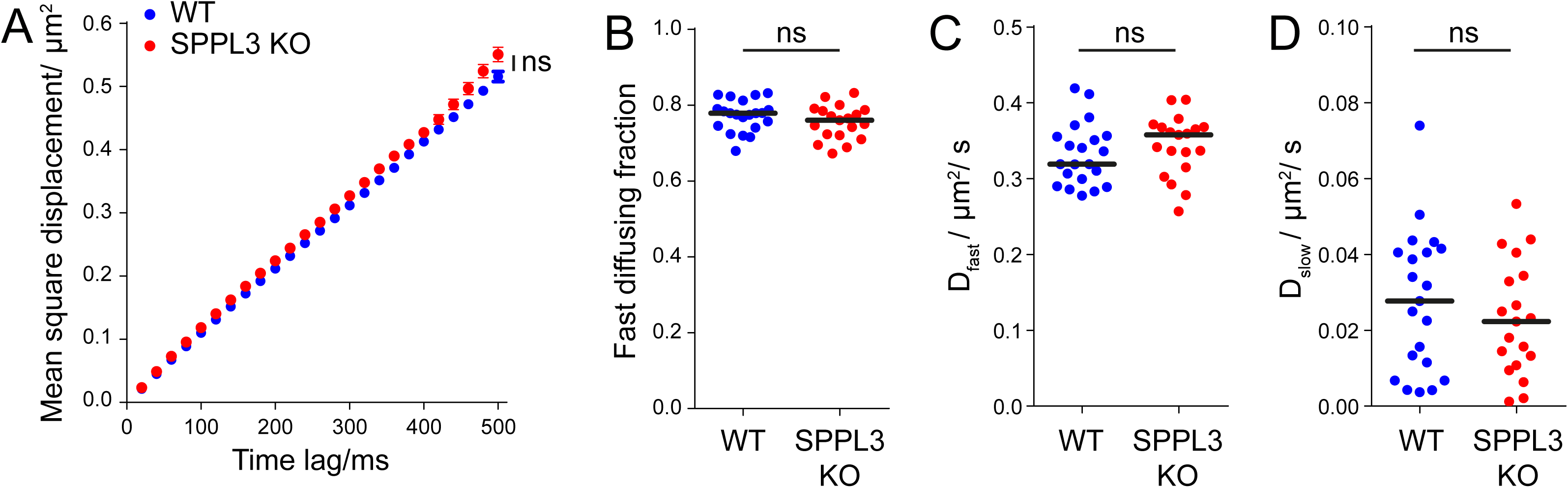
HLA-I membrane dynamics are not affected in the absence of SPPL3. Single particle tracking of individual HLA-I molecules was performed on WT and SPPL3 KO cells using SPPL3- independent HLA-I specific Alexa Fluor 555-conjugated BB7.2 Fab fragments. Single HLA-I molecules were tracked at room temperature with a time lag of 20ms. Representative example of n=3 is shown. (A) Mean square displacement plot of recorded HLA-I trajectories (mean and standard deviation are shown). Relative fractions (B) and diffusion constants of fast- (C) and slow-diffusing (D) HLA-I molecules on WT (n=21) or SPPL3 KO (n=20) cells were calculated by fitting the recorded HLA-I trajectories to a binary diffusion model representing a fast and slowly moving fraction. Statistics: median; Mann-Whitney U test.

**Figure S7.**
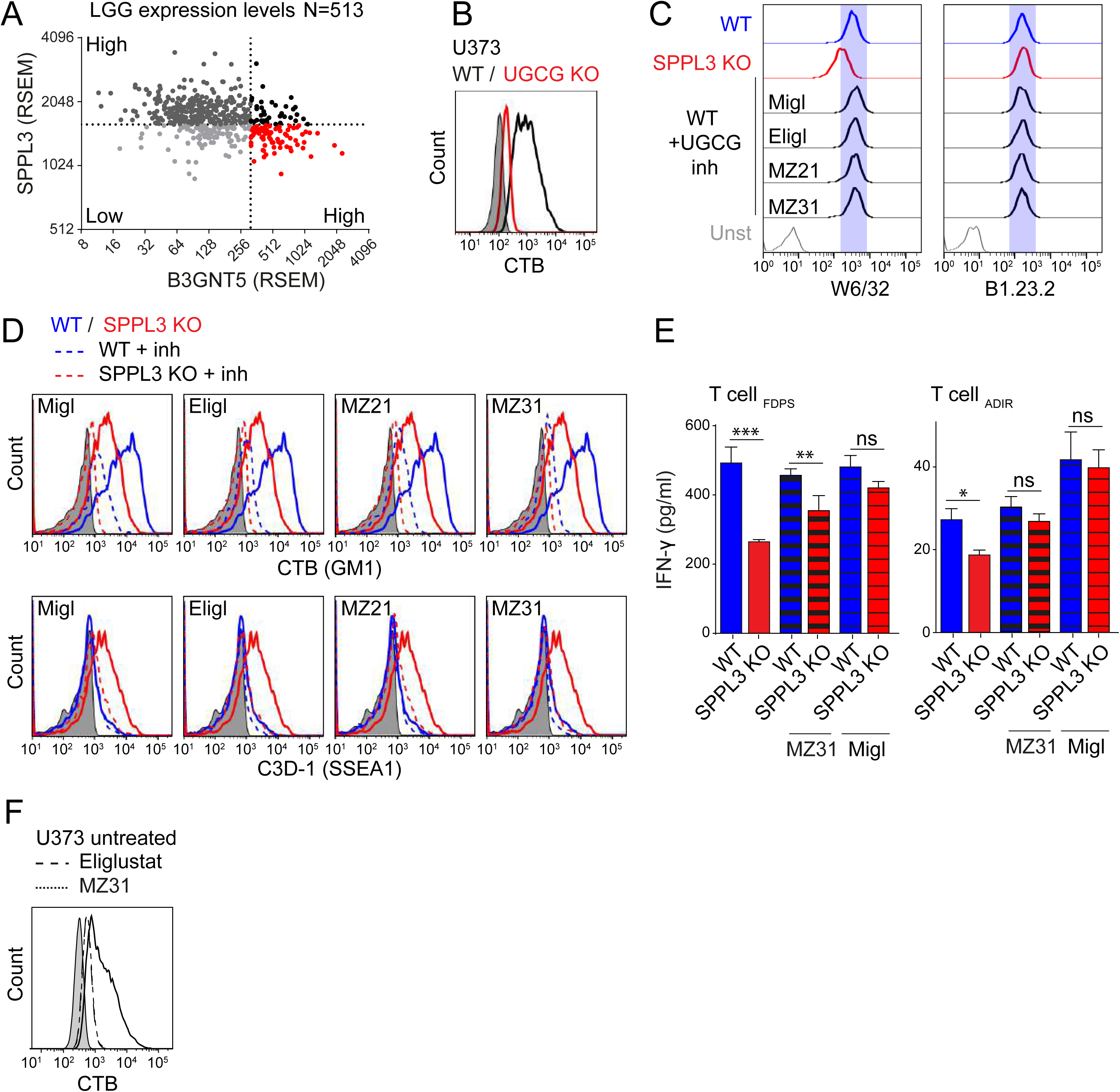
**Pharmacological inhibition of GSL synthesis in glioma enhances anti-tumor immune responses (*Relates to*** Figure 7) (A) Expression distribution of SPPL3 versus B3GNT5 in Low Grade Glioma (LGG) derived from TCGA. (B) Representative histograms of CTB cell surface staining of WT (*black*) and UGCG KO (*red*) U373 cells. Unstained control is in gray, n=2. (C)(D) Representative histograms of (C) W6/32 (*left*) and B1.23.2 (*right*), and (D) Cholera Toxin B stain (GM1) and C3D-1 (SSEA-1) cell surface staining of HAP1 WT (*blue*), SPPL3 KO (*red*) and WT cells preincubated with UGCG inhibitors miglustat, eliglustat, MZ21 and MZ31 (*black (in C); dashed (in D)*). Unstained control is in gray. The blue reference area indicates fluorescence of WT cells. n=3. (E) IFN- γ secretion by HLA-A*0201-restricted T cells recognizing FDPS (n=2), or ADIR (n=2) antigens on HAP1 WT (*blue*) or SPPL3 KO cells (*red*) each pre-incubated with (*dashed*) or without (*solid*) the indicated UGCG inhibitor (MZ31 or miglustat) as determined by ELISA. Representative plots of triplicate cultures are shown. (F) Representative histogram of CTB cell surface staining of WT (*solid*) U373 cells preincubated with UGCG inhibitors eliglustat (dashed) or MZ31 (*dotted*). Unstained control is in gray. n=3.

## EXTENDED MATERIALS AND METHODS

### Cell culture

HAP1 (HLA-A*02:01, HLA-B*40:01 and HLA-Cw*03:04), MelJuSo (HLA-A*01:01, B*08:01 and C*07:01), SW620 (HLA-A*24:02, A*02:01, B*07, B*15 and C*07:04) (kindly provided by Dr. T. de Gruijl (Amsterdam UMC, The Netherlands)) and U373 (HLA-A*02:01) (kindly provided by Dr. H. Versteeg (LUMC)) cell lines were cultured in IMDM (Gibco and Lonza) supplemented with 10% FCS and antibiotics (PenStrep; Invitrogen) at 37°C and 5% CO2. HEK293T and Phoenix ampho cells were cultured in similarly supplemented DMEM (Gibco). The HLA-A*0201-restricted CD8^+^ T cell clones reactive against peptides derived from the tumor-expressed proteins USP11, FDPS, VPS13B, ADIR, and SSR1 were previously described (Amir et al., 2011; van Bergen et al., 2007; Van Bergen et al., 2010) and expanded using a feeder cell-cytokine mixture in medium with human serum (Sanquin) as described (Oostvogels et al., 2014). Experiments with human materials were approved by the local medical ethical committee.

### Antibodies and LIR-1 Fc fusion protein

*(<u>Flow cytometry</u>)* W6/32-PerCP-eFluor710 (HLA-A, -B and -C), B1.23.2-APC (HLA-B and –C) and BB7.2-APC (HLA-A2) (eBioscience) were used to stain HLA-I. Additional HLA-I specific antibodies (all hybridoma supernatants, some purified using protein A) and their critical amino acids for their binding (epitope) are listed in Table S3. C3D-1-FITC (Millipore) and CTB-FITC (Sigma) were used to stain for respectively SSEA-1 and GM1. Goat anti-mouse IgG (H+L) AF647 (Invitrogen), mouse anti-human IgM (MHM-88, Biolegend) and mouse anti-human IgG (MH161-1, Sanquin) in-house conjugated to DL650 (Thermo Fisher Scientific) were used as secondary antibodies. The LIR-1 Fc fusion protein (human IgG1) was kindly provided by Dr. O. Mandelboim (Hebrew University Hadassah Medical School, Israel) (Gonen-Gross et al., 2010) and obtained from R&D Systems. (*<u>Single particle tracking</u>*) W6/32 Fab fragments were prepared by 1h pepsin (1μg/μL, pH3.5) treatment at 37°C in a pH3.5 buffer containing citric acid (0.07M) and sodium citrate (0.03M), followed by reduction using DTT (2.5mM, Sigma-Aldrich). BB7.2 Fab fragments were prepared using a papain:antibody ratio of 1:100 (16μg/mL papain) for 20h at 37°C. All Fab fragments were purified by gel filtration (Superdex 200, 10/300 GL, GE Healthcare Life Sciences). Monomeric Fab fragments were either conjugated to Alexa Fluor® 555 NHS Ester (AF555; ThermoFisher Scientific) or Alexa Fluor® 647 NHS Ester (AF647; ThermoFisher Scientific) according to the manufacturer’s labeling protocol. To remove unconjugated fluorophores, the labeled Fab fragments were further purified by gel filtration (Superdex 75, 10/300 GL, GE Healthcare Life Sciences). Fractions containing monomeric fluorophore-conjugated Fab fragments were concentrated to a protein concentration of 0.2-0.5mg/mL using Amicon Ultra-4 centrifugal filters (10 kDa cut off, Merck Millipore) and then stored in 50% glycerol at -20°C. The protein to dye ratios were determined by spectrophotometry at 280nm and the corresponding absorption maximum of the dyes at 555nm and 650nm. The protein to dye ratio of the randomly conjugated Fab fragments were 1.0 (BB7.2 Fab- AF555), 0.94 (W6/32 Fab-AF647) and 0.4 (W6/32 Fab-AF555). *(<u>Immunoprecipitation</u>)* W6/32 or BB7.2 antibodies (hybridoma supernatant) were used to immunoprecipitate HLA-I molecules. *(<u>Western blot</u>)* HC10/HCA2 (mixed hybridoma supernatants), anti-RFP (homemade), anti-FLAG M2 (Sigma) and anti-actin (AC15, Sigma) were used to stain HLA-I, RFP-tag, FLAG-tag, and actin respectively.

### Flow cytometry

Trypsinized cells were incubated with specific antibodies or proteins in PBS for 30min at 4°C. Samples stained with a secondary antibody were first washed three times in PBS/BSA. Stained cells were fixed in PBS containing 1% formaldehyde (Merck) and DAPI (1μM, Sigma-Aldrich) for direct analysis by flow cytometry. Samples were analyzed or sorted on BD flow cytometers (Canto, Fortessa, LSR II or ARIA II). FACS data was analyzed using FlowJo (Tree Star, Inc). Fluorescent cell barcoding was performed using the fluorescent dyes CFSE (125nM, Invitrogen), Alexa Fluor 350 NHS Ester (40μM, Thermo Fisher Scientific) and Violet proliferation dye 450 (2.5μM, BD Horizon). Pellet of cells was resuspendend in PBS containing the fluorescent dye and vortexed after every 5min for 15min. Barcoding was stopped by washing the cells three times in an excess ice-cold IMDM (Gibco and Lonza) supplemented with 10% FCS and antibiotics (PenStrep; Invitrogen). Barcoded cells were mixed prior to plating in 96V-bottom wells for antibody staining.

### Haploid genetic screening

Genome-wide knockout screening was performed in either early passage WT or CRISPR/Cas9 generated SPPL3 KO HAP1 cells using directly conjugated W6/32 or BB7.2 antibodies. Retroviral mutagenesis was performed on ∼100 x 10^6^ cells using GT-GFP or GT- BFP plasmids as previously described (Brockmann et al., 2017). In brief, 2 x 10^9^ expanded mutagenized cells were fixed, stained and sorted into two separate populations based on the fluorescent intensity of the respective HLA-I antibody staining (Brockmann et al., 2017). Gene trap integration sites were amplified using a LAM-PCR with a biotinylated primer on genomic DNA isolated from sorted cells. Biotinylated products were captured on streptavidin-coated beads followed by a single-stranded DNA linker ligation and a subsequent amplification step using two primers to generate fragments that include a genomic region flanking the insertion site in addition to adaptors for deep sequencing. Deep sequencing reads were aligned to the human genome (hg19) and intersected with protein coding genes to obtain the numbers of unique disruptive integrations mapped per gene in both populations either lowly or highly recognized by the respective HLA-I antibody. Enrichment of mutations in genes was assessed using a Fisher’s exact test corrected for false discovery (Benjamini-Hochberg). The approach is described in detail in (Brockmann et al., 2017).

### Immunoprecipitation

Transiently transfected HEK293T cells or CRISPR/Cas9 edited HAP1 cells (confluent 6cm dish) were lysed for 20min in lysis buffer containing 0.8% NP-40, 10% glycerol, 150mM NaCl, 50mM Tris-HCl pH8.0, 1mM EDTA, 5mM MgCl2 and protease inhibitors (Roche Diagnostics, EDTA free). Supernatant was centrifuged for 20min at 12.000g and incubated with RFP- Trap beads (Chromotek) or antibody coated Protein-G sepharose beads for 2h. Beads were washed four times in lysis buffer before addition of Laemmli Sample Buffer (containing 5% β-mercaptoethanol) followed by 5min incubation at 95°C. Co-immunoprecipitated proteins were separated by SDS-PAGE for Western blotting and detection by antibody staining. N-glycosylation inhibitor activity was confirmed by Endo H treatment of immunoprecipitated HLA-I molecules (using W6/32 hybridoma supernatant) in 50nM Nacitrate (pH5.5)/0.2% SDS according to manufacturer’s protocol (Sigma). HLA-I molecules used for glycopeptide analysis were immunoprecipitated using W6/32.

### SDS-PAGE and Western blotting

Samples were separated by SDS-PAGE (10% acrylamide gel) and transferred to a PVDF membrane (Immobilon-P, 0.45μm, Millipore) at 300mA for 3h. The filters were blocked in PBS/5% Milk (Skim milk powder, Oxoid) and incubated with a primary antibody for 1h diluted in PBS/0.1% Tween/5% Milk, washed thrice for 10min in PBS/0.1% Tween and incubated with the secondary antibody for 45min diluted in PBS/0.1% Tween/5% Milk and washed thrice again in PBS/0.1% Tween. The filter was incubated with ECL reagent (SuperSignal West Dura Extended Duration Substrate, Thermo Fisher Scientific) and the signal was detected using the Chemidoc XRS+ imager (Bio-Rad) or Amersham Imager 600.

### Inhibitors and enzymes

UGCG inhibitors MZ31 (use concentration: 2µM), MZ21 (2µM), miglustat (100µM) were produced as previously described (Ghisaidoobe et al., 2014) and eliglustat (200nM) was obtained from Bio-Connect. Swainsonine (20µg/mL) and kifunensine (25µM) were obtained from Sigma. Sialyltransferase inhibitor (3Fax-peracetyl Neu5Ac, 100µM) was obtained from Sigma and fucosyltransferase inhibitor (2-Deoxy-2-fluoro-L-fucose, 100µM) was obtained from Carbosynth. Cells were incubated for 2 days at 37 degrees with inhibitors before flow cytometry analysis. Cells were incubated with Neuraminidase (N2876, Sigma, 225mU/ml) for 1h at 37 degrees.

### Overexpression and genome editing

pMXs-puro vector (Cell Biolabs) was equipped with a novel multiple cloning site with or without N- or C-terminal tags (RFP and FLAG). SPPL3 and its inactive mutant SPPL3 D271A (kind gift from Dr. R. Flührer) (Voss et al., 2012) were recloned into pMXs-puro/RFP-N1 using XhoI/BamHI restriction sites. B3GNT5 and B4GALNT1 were PCR amplified from IMAGE:202800754 and IMAGE:202800771 and cloned into pMXs-puro/FLAG-C1 by EcoRI/BamHI restriction sites. ST3GAL5 was amplified from IMAGE:202759803 and cloned into pMXs-puro/FLAG- C1 using EcoRI/BclI digestion into an EcoRI/BamHI digested plasmid. The pLZRS-based retroviral vectors containing HLA-A*02:01/B*40:01/C*03:03-IRES-ΔNGFR were described before (Griffioen et al., 2012; Van Bergen et al., 2010). Generation of retroviral supernatants and transduction of cells were performed as described (Spaapen et al., 2008). For genome editing, gRNAs (Table S4) were cloned into the pX330 expression vector or the lentiviral vectors lentiCRISPR_v2 or pLCRISPR.efs.GFP (Addgene) (Heckl et al., 2014; Sanjana et al., 2014). SPPL3 KO and tapasin KO HAP1 cells were created by in frame integration of a blasticidin-resistance gene after cotransfection of pX330 with TIA-2Ablast (using Extremegene HP, Sigma) as described for other targets (Blomen et al., 2015; de Waard, 2020). Clones growing after blasticidin selection (10µg/mL, Life Technologies) were sequence verified for specific genome editing by Sanger sequencing (primers in Table S5). Lentiviral constructs containing gRNAs targeting SPPL3, UGCG, CMAS and the five core GSL-enzymes were co-transfected into HEK293T with the packaging enzymes psPAX2, pVSVg, pAdVAntage using polyethylenimine (PEI; Polyscience) for virus production. Filtered viral supernatants were used for transduction by spinoculation in the presence of 8µg/mL protamine sulfate. Cells were selected using puromycin (0.25µg/mL; Gibco), blasticidin (10µg/mL; Gibco), or gated on based on the co-expression of GFP. Polyclonal KO populations after selection were used for flow cytometry (SPPL3, CMAS and core GSL-enzymes) or KO clones were made by limiting dilution and sequence verified. HLA-A, -B and -C KO cells were generated by pX330 transfection followed by single cell FACS sort using W6/32 and sequence verification of clones (de Waard, 2020).

### Sanger sequencing

Constructs were sequenced using BigDye Terminator Kit (Applied Biosystems). Genomic DNA isolated from cell lines using DirectPCR (Cell) lysis reagent (Viagen) supplemented with Proteinase K (Sigma) was amplified using primers mentioned in Table S5 and directly sequenced using BigDye Terminator Kit (Applied Biosystems). Sequences were analyzed using Snapgene (GSL Biotech). Sequence decomposition to assess the size of deletions or insertions of genome-edited diploid cells was performed using Tide (Brinkman et al., 2014).

### siRNA transfections

Gene silencing was performed in a 96F well plate using 5μL siRNA (500nM stock) mixed with 0.1μL DharmaFECT1 #1 (Dharmacon) in 4.9μL IMDM similar as we described before (Paul et al., 2011). After three days cells were analyzed using FACS. SPPL3 and B2M were silenced using siRNAs (Dharmacon). Non-targeting siRNA (siCTRL, D-001206-13-20, Dharmacon) was used as a negative control.

### Preparation of fibronectin-coated glass slides for single particle tracking experiments

Glass slides (24mm x 50mm #1.5 borosilicate, VWR) were immersed in a 1:1 mixture of concentrated sulfuric acid (Sigma) and 30% hydrogen peroxide (Sigma) for at least 30min, rinsed with deionized water, air dried and glued with picodent twinsil extrahart (Picodent) to the bottom of 8-well LabTek chambers (Nunc). Slides were coated with 20µg/ml fibronectin (Sigma-Aldrich) in PBS for 1-2h at 37°C and rinsed with 1 X PBS.

### Microscopy

We employed an Eclipse Ti-E (Nikon) inverted microscope system to image samples in epifluorescence or total internal reflection fluorescence mode (TIRF) with a high NA objective (100X magnification, NA=1.49, Nikon SR APO TIRF). When indicated images were obtained using 20X magnification (NA=0.75, Nikon S Fluor). The microscope system was further equipped with five diode lasers featuring 405nm, 488nm, 514nm, 642nm (iBeam smart Toptica) and 532nm and 561nm (Obis) light. The laser excitation light was cleaned up with MaxLine clean up filters that matched the corresponding wavelengths (Semrock). The setup was equipped with the beam splitters zt405/488/561/647rpc, zt405/488/532/640rpc, and zt405/514/635rpc (AHF). To clean up the emission path, we used a fast filter wheel (Sutter Instrument Company) having the custom bandpass filters ET450/50, ET510/20, ET525/50, ET605/52 and ET700/75 (all AHF) installed. Emission light was recorded with a back-illuminated EM-CCD camera (Andor iXon Ultra 897). Temperature-sensitive experiments were conducted under the control of a temperature control system from Pecon. An 8 channel DAQ-card (National Instruments) and the microscopy automation and image analysis software Metamorph (Molecular Devices) were used to program and apply timing protocols and control all hardware components.

### Single particle tracking of HLA-I molecules on HAP1 WT and SPPL3 KO cells

0.2 x 10^6^ HAP1 WT and SPPL3 KO cells were stained with a single molecule dilution of the BB7.2-Fab-AF555 on ice for 30min and washed twice in imaging buffer (HBSS, Gibco, 1% FCS, 2mM MgCl2 and CaCl2). Cells were kept on ice or seeded onto fibronectin-coated glass slides for imaging at room temperature (23- 27°C) and in TIRF mode. AF555 was excited with a 532 nm laser (Obis) with a power density of 0.8kW/cm^2^ and the emission channel was cleaned up with a TRITC filter (605/52) installed within the fast emission filter wheel (Sutter Instrument Company). We recorded single HLA-I (BB7.2 Fab-AF555) trajectories over 500 frames with an illumination time of 16milliseconds and a total time lag of 20.6milliseconds between two adjacent images (100 X 100 pixel ROI size). Microscopy images were processed and analyzed with the open-source image processing package Fiji (Schindelin et al., 2012). XY localization, intensity and positional accuracy of single fluorescence emitters was calculated with the Fiji plugin ThunderSTORM (Ovesny et al., 2014). After determining the localization of every fluorophore in the image stack we combined these localizations to trajectories based on a published approach (Gao and Kilfoil, 2009) with custom-made algorithms written in Matlab (MathWorks). We calculated the mean square displacement (MSD) describing the average of the square displacements between two points of the trajectory according to *MSD(tlag) = <(r•(t + tlag)-r(t))²*>. The first three MSD values as a function of time lag (tlag) were used to calculate the diffusion coefficient (D) for each trajectory according by fitting *MSD=4•D•tlag+4•σxy²*, with *σxy* representing the localization precision (Wieser et al., 2007). Multiple fractions (i.e. a fast and a slow moving fraction of molecules) were discriminated by analyzing the step-size distributions of square displacements for several time lags (Schutz et al., 1997). By assuming free Brownian motion of one mobile fraction, the cumulative probability for finding a square displacement smaller than r² is given by 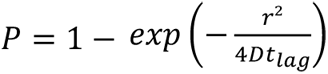; two different fractions *α* and *(1-α)* with diffusion coefficients D1 (e.g. fast) and D2 (e.g. slow/immobile) can be distinguished by fitting the bi-exponential function 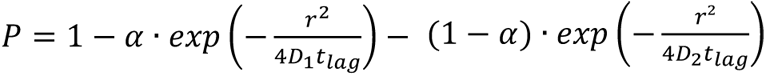. We calculated the fraction of mobile and slow/immobile molecules, the diffusion rate of mobile molecules and the diffusion rate of slow/immobile molecules of all HLA-I trajectories present on a single cell and summarized the data in a box plot (Brameshuber et al., 2018).

### B3GNT5 activity assay

5 x 10^6^ HAP1 cells were harvested by trypsinization and cells were lysed in lysis buffer (2% TritonX-100, 50mM sodium cacodylate pH7.4, 10mM MnCl2, and complete EDTA-free protease inhibitor cocktail (Roche) by incubating on ice for 30min. Nuclei were precipitated by centrifugation at 15,000x *g.* Next, post-nuclear supernatants containing equal amounts of protein in 50µL lysis buffer were added equal volume of lysis buffer containing 2µM BODIPY-C5- Lactosylceramide complexed to BSA (Thermo Fisher Scientific) and/or 1mM UDP-*N*-acetyl-D-glucosamine (Santa Cruz). Samples were incubated at 37°C for 4h and subjected to lipid extraction using Bligh-Dyer method (Bligh and Dyer, 1959; Jongsma et al., 2017). Briefly, reactions were added 100µL 2% NaCl, 250µL chloroform, and 500µL methanol and samples were vortexed. Phase separation was induced by addition of 250µl 0.45% NaCl and 250µL chloroform and lower phases were collected. Upper phases were re-extracted twice more and collected lower phases were dried under a nitrogen flow. Dried lipid were re-suspended in chloroform:methanol (2:1 v/v) solution and spotted on TLC plates. TLC plates were developed in chloroform:methanol:water (60:25:4 v/v/v) and imaged using Typhoon FLA9500 (GE Healthcare) scanner equipped with a 473nm laser and BPB1 filter (530DF30). Identity of BODIPY-Lc3Cer was confirmed by MS/MS. Structures were assigned based on MS/MS fragmentation pattern in negative mode following nomenclature from Domon and Costello (Domon and Costello, 1988).

### Glycosphingolipid extraction and purification by RP-SPE

GSLs were extracted from 1 x 10^7^ HAP1 WT, SPPL3 KO and B3GNT5/SPPL3 double KO cell lines in triplicate in glass vials equipped with a Teflon-lined screw cap. Cells were washed three times in 1mL of water followed by centrifugation at 2000xg for 30min. The supernatant was removed and replaced by 300μL of 2- propanol. The samples were vortexed for 5min and incubated for 15min at 75°C. A volume of 350μL of MTBE was added to the samples followed by 15min sonication. 200μL of water was added to the cell pellets and incubated for 4h with shaking at room temperature. The upper phase containing GSLs was collected after centrifugation at 2700xg for 20min. Then, 400μL of MTBE was added, followed by sonication and centrifugation. The upper phase was collected and pooled to the previous sample. The process of adding MTBE, sonication, centrifugation and removing upper phase was repeated for another two times. The combined upper phases were dried under vacuum in an Eppendorf Concentrator 5301 (Eppendorf) at 30°C. Before purification of the GSLs using RP-solid phase extraction (SPE), the samples were dissolved in 200μL methanol and vortexed for 10min, followed by addition of 400μL water. TC18-RP-catridges were prewashed with 2mL of chloroform/methanol (2:1, v/v), 2mL of methanol followed by equilibration with 2mL methanol/water (1:2, v/v). The extracted GSLs were loaded to the cartridge for 3 times and washed with 3mL methanol/water (1:2, v/v). The GSLs were eluted from the column with 2mL methanol and 2mL chloroform/methanol (2:1, v/v). The samples containing the eluate were evaporated under nitrogen for 1h and dried under vacuum in an Eppendorf Concentrator at 30°C. The collection and dry of GSLs eluate were performed in glass tube.

### GSL glycan release by EGCase I and purification

To release the glycans from the GSLs, a mixture of EGCase I (12mU, 2μL), EGCase I buffer (4μL) and water (34μL) (pH5.2) was added to each sample and incubated for 16h at 37°C. The released glycans were collected and applied to a TC18-RP-cartridges which was preconditioned with 2mL of methanol and 2mL of water. The sample vials were washed with 200µL of water and residual glycans were loaded to the cartridge. Then, 500μL of water was added to the cartridge to wash the glycans from the column. The flow-through and wash fractions were pooled and dried in an Eppendorf Concentrator at 30°C.

### Reduction, desalting and carbon SPE cleanup of GSL glycans

The reduction was carried out as described previously with slight modifications (Jensen et al., 2012). In brief, GSL glycans were reduced to alditols in 20μL of sodium borohydride (500mM) in potassium hydroxide (50mM) for 2h at 50°C. Subsequently, 2μL of 100% glacial acetic acid was added to neutralize the solution and quench the reaction. The desalting of GSL glycans was performed on cation exchange columns which consist of 60μL of AG50W-X8 resin beads deposited onto reversed phase μC18 ZipTips (Perfect Pure, Millipore) as previously described (Jensen et al., 2012). Glycan alditols were eluted with 50μL of water twice. The combined flow-through and eluate were pooled and dried under vacuum in an Eppendorf Concentrator at 30°C. The carbon SPE clean-up was performed and the purified glycan alditols were resuspended in 10μL water for porous graphitized carbon (PGC) LC-ESI-MS/MS analysis.

### Analysis of GSL glycans using PGC LC-ESI-MS/MS

Porous graphitized carbon (PGC) LC-ESI-MS/MS analysis of GSL glycan alditols was performed on a Dionex Ultimate 3000 nano-LC system equipped with a Hypercarb PGC trap column (5μm Hypercarb Kappa, 32μm x 30mm, Thermo Fisher Scientific) and a Hypercarb PGC nano-column (3μm Hypercarb Kappa, 75μm x 100mm, Thermo Fisher Scientific) coupled to an amaZon speed ion trap mass spectrometer (Bruker Daltonics). Mobile phase A consisted of 10mM ammonium bicarbonate. Mobile phase B was 60% (v/v) acetonitrile / 10mM ammonium bicarbonate. To analyze glycans, 2μL injections were performed and separation was achieved with a gradient of B (1-71% at 0.7%/min) followed by a 10min wash step using 95% of B at a flow of rate of 0.6μL/min. MS scans from *m/z* 340 to 1700 were recorded in enhanced mode using negative ion mode. MS/MS spectra were recorded selecting the top 3 highest intensity peaks. Glycan structures were assigned based on glycan composition obtained from accurate mass, relative PGC elution position, MS/MS fragmentation pattern in negative-ion mode and general glycobiological knowledge (Karlsson et al., 2004), with help of Glycoworkbench (Ceroni et al., 2008) and Glycomod (Cooper et al., 2001) software tools. Extracted ion chromatograms were used to integrate area under the curve (AUC) for each individual glycan isomer using Compass Data Analysis software v.5.0. The most abundant peaks in the glycan profile were manually picked and integrated. Relative quantitation of individual glycans was performed on the total area of all included glycans within one sample normalizing it to 100%.

### In-gel tryptic digestion

The glycopeptide generation and analysis were performed as described previously with slight modifications (Plomp et al., 2014). BB7.2 immunoprecipitated samples were loaded on SDS PAGE. Bands containing HLA-I were excised and cut into pieces. The gel pieces were washed with 25mM ammonium bicarbonate and dehydrated with acetonitrile. The samples were then reduced in-gel for 30min at 55°C with 100μL 10mM DTT in 25mM ammonium bicarbonate solution, followed by dehydration of the gel pieces in ACN and cysteine alkylation for 20min with 100μL of a 55mM iodoacetamide 25mM ammonium bicarbonate solution in the dark for 45min. The gel pieces were washed in 25mM ammonium bicarbonate, followed by removal of the solution and incubation in ACN. This was repeated twice, and the gel pieces were subsequently dried in a centrifugal vacuum concentrator at 30°C for 10min. Enzymatic digestion of trypsin was performed by adding 50μL of 25mM ammonium bicarbonate containing 0.6μg of trypsin (sequencing grade modified trypsin, Promega) to the dried gel particles. The samples were kept on ice for 1 h and were subsequently incubated overnight at 37°C. The solution surrounding the gel pieces was collected and stored at - 20°C. We then added 20μL of 25mM ammonium bicarbonate to the gel pieces, and incubation was continued at 37°C degrees for another hour. The solution was again collected and added to the first fraction prior to freezing.

### Glycopeptide analysis by LC-ESI-MS/MS

Glycopeptides were analyzed by nano-RP-LC-ESI-ion trap-MS(/MS) on an Ultimate 3000 RSLCnano system (Dionex / Thermo Fisher Scientific) coupled to an HCTultra-ESI−IT-MS (Bruker Daltonics). Five microliters of sample was injected and concentrated on a trap column (Acclaim PepMap100 C18 column, 100μm × 2cm, C18 particle size 5μm, pore size 100Å, Dionex / Thermo Fisher Scientific) before separation on an Acclaim PepMap RSLC nanocolumn (75μm × 15cm, C18 particle size 2μm, pore size 100Å, Dionex / Thermo Fisher Scientific). A flow rate of 700nL/min was applied. Solvent A consisted of 0.1% formic acid in water; solvent B, 0.1% formic acid in 95% ACN and 5% water. A linear gradient was applied with the following conditions: *t* = 0min, 3% solvent B; *t* = 5min, 3% solvent B; *t* = 20min, 27% solvent B; *t* = 21min, 70% solvent B; *t* = 23min, 70% solvent B; *t* = 24min, 3% solvent B; *t* = 43min, 3% solvent B. Samples were ionized in positive ion mode with an online nanospray source (4500V) using fused- silica capillaries and a Distal Coated SilicaTip Emitter (New Objective) with an internal diameter of 20μm (10μm at the tip) and a length of 5cm. Solvent evaporation was performed at 220°C with a nitrogen flow of 3L/min. For the detection of glycopeptides, the MS ion detection window was set at *m/z* 500-1800, and the MS/MS detection window at *m/z* 140-2200, with automated selection of the three highest peaks in the spectrum for MS/MS analysis. The LC-MS/MS results were analyzed using DataAnalysis 4.0 software (Bruker Daltonics) and screened manually for the masses of common oxonium fragment ions (*m/z* 366.1, [1 hexose + 1 GlcNAc + H]^+^ ; *m/z* 657.2, [1 hexose + 1 GlcNAc + 1 *N*-acetylneuraminic acid + H]^+^ ; *m/z* 528.2, [2 hexoses + 1 GlcNAc + H]^+^), which are characteristic for fragmentation spectra of glycopeptides. Glycopeptide MS/MS spectra were further analyzed manually to derive the oligosaccharide structure and the mass of the peptide moiety.

### Other materials

Iodoacetamide was obtained from Sigma-Aldrich. bicarbonate, cation exchange resin beads (AG50W-X8), methyl tert-butyl ether (MTBE), trifluoroacetic acid (TFA), potassium hydroxide, methanol, ammonium bicarbonate and sodium borohydride were obtained from Sigma- Aldrich. Endoglycoceramidase I (EGCase I, recombinant clone derived from *Rhodococcus triatomea* and expressed in *Escherichia coli*) and 10x EGCase I buffer (500mM HEPES, 1M NaCl, 20mM DTT and 0.1% Brij 35, pH5.2) were purchased from New England BioLabs. HPLC SupraGradient acetonitrile (ACN) was obtained from Biosolve and other reagents and solvents such as methanol, ethanol, chloroform, 2-propanol, and glacial acetic acid were from Merck. The 50mg TC18-reverse phase (RP)-cartridges were from Waters. Ultrapure water was used for the all preparations and washes, generated from a Q-Gard 2 system (Millipore).

### Crystal structures

Structural prediction software Phyre2 was used to create a model of SPPL3 using the primary consensus sequence CCDS9208.1 (Kelley et al., 2015). Models of HLA-A2 were made using the crystal structure 3MRG courtesy of the RCSB PDB (Reiser et al., 2014; Winn et al., 2011). The structure 3QZW was used in conjunction with the CCP4 programme ARIAIMOL (Berman et al., 2000) to determine the hydrogen bonding contacts between human HLA-A*2402 and human CD8 alpha-alpha dimer (Shi et al., 2011). A similar method was used to assess the contacts between LIR-1 and HLA-A2 using the structure 1P7Q (Berman et al., 2000; Willcox et al., 2003). All figures have been produced using the PyMOL molecular graphics software (Version 2.0 Schrödinger, LLC).

### In silico and statistical analyses

All error bars correspond to the standard deviation of the mean. Data from genome-wide screens were analyzed using two-sided Fisher’s exact test followed by FDR (Benjamini-Hochberg) correction of the p-value. Other statistical evaluations were done by a Student’s t-test (analysis of two data groups), one-way ANOVA (three groups or more), two-way ANOVA (two variables), Mann-Whitney U test (non-parametric analyses) or Log-rank (survival analyses) with Prism software (http://www.graphpad.com). TCGA survival and expression data were retrieved from OncoLnc.org (Anaya, 2016). *p<0.05, **p<0.01, ***p<0.001 and ns (= not significant). EC50 values of titrations were calculated using non-linear four parameter fit modeling with Prism software.

